# PICH function is required for organization of SUMOylated proteins on mitotic chromosomes

**DOI:** 10.1101/2020.02.06.937243

**Authors:** Victoria A. Hassebroek, Hyewon Park, Nootan Pandey, Brooklyn T. Lerbakken, Vasilisa Aksenova, Alexei Arnaoutov, Mary Dasso, Yoshiaki Azuma

**Affiliations:** Department of Molecular Biosciences, University of Kansas, Lawrence, Kansas, U.S.A, 66045; Division of Molecular and Cellular Biology, National Institute for Child Health and Human Development, National Institutes of Health, Bethesda, MD 20892, USA

**Author notes:** To whom correspondence should be addressed: Yoshiaki Azuma: Department of Molecular Biosciences, University of Kansas, Lawrence, Kansas, U.S.A, 66045.

**Keywords:** Chromosome, Mitosis, PICH, SUMO, TopoisomeraseIIα

## Abstract

Proper chromosome segregation is essential for faithful cell division and if not maintained results in defective cell function caused by abnormal distribution of genetic information. Polo-like kinase 1 interacting checkpoint helicase (PICH) is a DNA translocase essential in chromosome bridge resolution during mitosis. Its function in resolving chromosome bridges requires both DNA translocase activity and ability to bind chromosomal proteins modified by Small Ubiquitin-like modifier (SUMO). However, it is unclear how these activities are cooperating to resolve chromosome bridges. Here, we show that PICH specifically promotes the organization of SUMOylated proteins like SUMOylated TopoisomeraseIIα (TopoIIα) on mitotic chromosomes. Conditional depletion of PICH using the Auxin Inducible Degron (AID) system resulted in the retention of SUMOylated chromosomal proteins, including TopoIIα, indicating that PICH functions to control proper association of these proteins with chromosomes. Replacement of PICH with its mutants showed that PICH is required for the proper organization of SUMOylated proteins on chromosomes. *In vitro* assays showed that PICH specifically attenuates SUMOylated TopoIIα activity using its SUMO-binding ability. Taken together, we propose a novel function of PICH in remodeling SUMOylated proteins to ensure faithful chromosome segregation.

**Summary Statement:** Polo-like kinase interacting checkpoint helicase (PICH) interacts with SUMOylated proteins to mediate proper chromosome segregation during mitosis. The results demonstrate that PICH controls association of SUMOylated chromosomal proteins, including Topoisomerase IIα, and that function requires PICH translocase activity and SUMO binding ability.

## Introduction

Accurate chromosome segregation is a complex and highly regulated process during mitosis. Sister chromatid cohesion is necessary for proper chromosome alignment, and is mediated by both Cohesin and catenated DNA at centromeric regions (Michaelis *et al.*, 1997; Losada *et al.*, 1998; Bauer *et al.*, 2012). Compared to the well-described regulation of Cohesin (Morales and Losada, 2018), the regulation of catenated DNA cleavage by DNA TopoisomeraseIIα (TopoIIα) is not fully understood despite its critical role in chromosome segregation. ATP-dependent DNA decatenation by TopoIIα takes place during the metaphase-to-anaphase transition and this allows for proper chromosome segregation (Shamu and Murray, 1992; Wang *et al.*, 2010; Gomez *et al.*, 2014). Failure in resolution of catenanes by TopoIIα leads to the formation of chromosome bridges, and ultra-fine DNA bridges (UFBs) to which PICH localizes (Spence *et al.*, 2007). PICH is a SNF2 family DNA translocase (Baumann *et al.*, 2007; Biebricher *et al.*, 2013), and its binding to UFBs recruits other proteins to UFBs (Chan *et al.*, 2007; Hengeveld *et al.*, 2015). In addition to the role in UFB binding during anaphase, PICH has been shown to play a key role in chromosome segregation at the metaphase to anaphase transition (Baumann *et al.*, 2007; Nielsen *et al.*, 2015; Sridharan and Azuma, 2016).

Previously, we demonstrated that PICH binds SUMOylated proteins using its three SUMO interacting motifs (SIMs) (Sridharan *et al.*, 2015). PICH utilizes its ATPase activity to translocate DNA, similar to known nucleosome remodeling enzymes (Whitehouse *et al.*, 2003), thus it is a putative remodeling enzyme for chromosomal proteins. But, the nucleosome remodeling activity of PICH was shown to be limited as compared to established nucleosome remodeling factors (Ke *et al.*, 2011). Therefore, the target of PICH remodeling activity has not yet been determined. Importantly, both loss of function PICH mutants in either SUMO-binding activity or translocase activity showed chromosome bridge formation (Sridharan and Azuma, 2016), suggesting that both of these activities cooperate to accomplish proper chromosome segregation albeit the molecular mechanism linking these two functions is unknown. Previous studies demonstrated that proper regulation of mitotic chromosomal SUMOylation is required for faithful chromosome segregation (Nacerddine *et al.*, 2005; Diaz-Martinez *et al.*, 2006; Cubenas-Potts *et al.*, 2013). Studies using C. elegans demonstrated the dynamic nature of SUMOylated proteins during mitosis and its critical role in chromosome segregation (Pelisch *et al.*, 2014). Several SUMOylated chromosomal proteins were identified for their potential role in chromosome segregation, for example; TopoIIα, CENP-A, CENP-E, FoxM1, and Orc2 (Bachant *et al.*, 2002; Zhang *et al.*, 2008; Schimmel *et al.*, 2014; Huang *et al.*, 2016; Ohkuni *et al.*, 2018). Because PICH is able to specifically interact with SUMO moieties (Sridharan *et al.*, 2015), these SUMOylated chromosomal proteins could be a target of the SIM-dependent function of PICH in mediating faithful chromosome segregation. Among the known SUMOylated chromosomal proteins, TopoIIα has been shown to functionally interact with PICH. PICH-knockout cells have increased sensitivity to ICRF-193, a potent TopoII catalytic inhibitor, accompanied with increased incidence of chromosome bridges, binucleation, and micronuclei formation (Wang *et al.*, 2008; Kurasawa and Yu-Lee, 2010; Nielsen *et al.*, 2015). ICRF-193 stalls TopoIIα at the last step of the strand passage reaction (SPR) in which two DNA strands are trapped within the TopoIIα molecule without DNA strand breaks (Roca *et al.*, 1994; Patel *et al.*, 2000). In addition to that specific mode of inhibition, ICRF-193 has been shown to increase SUMOylation of TopoIIα (Agostinho *et al.*, 2008; Pandey *et al.*, 2020). Because PICH has SUMO binding ability, it is possible that increased SUMOylation of TopoIIα contributes to interaction with PICH under ICRF-193 treatment. However, no study has shown a linkage between SUMOylation of TopoIIα and PICH function.

To elucidate possible functional interactions of PICH with SUMOylated chromosomal proteins, we established the connection between PICH and SUMOylation by utilizing specific TopoII inhibitors and genome edited cell lines. Our results demonstrate that increased SUMOylation by ICRF-193 treatment leads to the recruitment of and enrichment of PICH on chromosomes. Depletion of SUMOylation abrogates this enrichment, suggesting PICH specifically targets SUMOylated chromosomal proteins. Depletion of PICH led to the retention of SUMOylated proteins including SUMOylated TopoIIα on the chromosomes in ICRF-193 treated cells. Replacing endogenous PICH with a translocase deficient PICH mutant resulted in increased SUMO2/3 foci on chromosomes where PICH was located, suggesting that PICH utilizes its translocase activity to remodel SUMOylated proteins on the chromosomes. *In vitro* assays showed that PICH specifically interacts with SUMOylated TopoIIα to attenuate SUMOylated TopoIIα activity in a SIM dependent manner. Together, we propose a novel mechanism for PICH in promoting proper chromosome segregation during mitosis by remodeling SUMOylated proteins on mitotic chromosomes including TopoIIα.

## Results

### Upregulation of SUMO2/3 modification by treatment with TopoII inhibitor ICRF-193 causes increased PICH foci on mitotic chromosomes

We previously reported that PICH utilized its SIMs for proper chromosome segregation and for its mitotic chromosomal localization (Sridharan and Azuma, 2016). We wished to examine whether modulating mitotic SUMOylation affected PICH localization on mitotic chromosomes. Treatment with ICRF-193, a catalytic inhibitor of TopoII which blocks TopoII at the last stage of its SPR, after DNA decatenation but before DNA release, increases SUMO2/3 modification of TopoIIα on mitotic chromosomes. In contrast, treatment with another catalytic TopoII inhibitor, Merbarone, which blocks TopoII before the cleavage step of the SPR, does not affect the level of SUMO2/3 modification of TopoIIα (Agostinho *et al.*, 2008; Pandey *et al.*, 2020). We utilized these two contrasting inhibitors to assess whether TopoIIα inhibition and/or SUMOylation changes PICH distribution on mitotic chromosomes. DLD-1 cells were synchronized in prometaphase, and mitotic cells were collected by mitotic shake off then chromosomes were isolated. To assess the effects of the TopoII inhibitors specifically during mitosis, the inhibitors were added to cells after mitotic shake off. Consistent with previous reports (Agostinho *et al.*, 2008; Pandey *et al.*, 2020), Western blot analysis of isolated chromosomes showed that treatment with ICRF-193 significantly increased the overall SUMO2/3 modification of chromosomal proteins including SUMOylated TopoIIα (marked by red asterisks in Figure 1A). Intriguingly, a novel finding showed that PICH levels on mitotic chromosomes were significantly increased in cells treated with ICRF-193. In contrast, Merbarone did not increase the level of these proteins on the chromosomes suggesting that there is a specificity of ICRF-193 which causes increased levels of PICH and SUMOylation of TopoIIα (Figure 1A).

**Figure 1.**
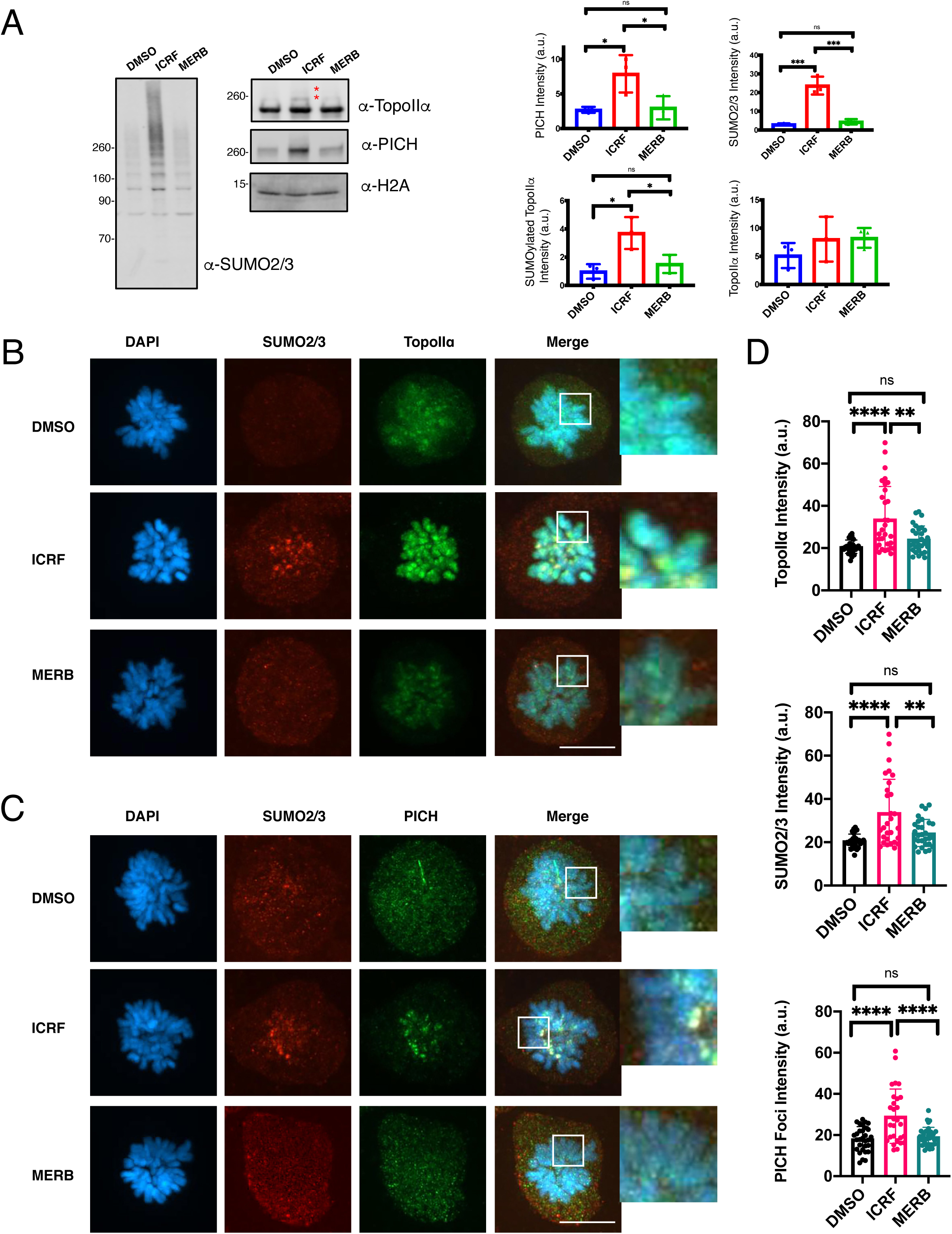
TopoIIα inhibition by ICRF-193 leads to increased PICH, SUMO2/3 and TopoIIα levels on mitotic chromosomes. **(A)** DLD-1 cells were synchronized and treated with indicated inhibitors (7μM ICRF-193: ICRF, and 40μM Merbarone: Merb), DMSO was used as a control. Mitotic chromosomes were isolated and subjected to Western blotting with indicated antibodies. Intensity of signals (arbitrary unit; a.u.) normalized by H2A are shown with mean value and standard deviation. * indicates SUMOylated TopoIIα. p values for comparison from three experiments were calculated using a one-way ANOVA analysis of variance with Tukey multi-comparison correction. ns: not significant; *: p ≤ 0.05; ***: p < 0.001 **(B)** Mitotic cells treated with DMSO (control), ICRF-193, and Merbarone were stained with antibodies against: TopoIIα (green) and SUMO2/3 (red). DNA was stained with DAPI (blue). Scale bar = 11μm. The white square indicates enlarged area. **(C)** Mitotic cells were treated as in **B** and stained with antibodies against: PICH (green), SUMO2/3 (red). DNA was stained with DAPI (blue). Scale bar = 11μm. The white square indicates enlarged area. **(D)** Using DAPI signal the mean intensity (a.u.) of each channel of at least 5 individual chromosomes per experimental replicate were measured. The bar indicates the mean value of the intensities. p values for comparison of all obtained values from three experiments were calculated using a one-way ANOVA analysis of variance with Tukey multi-comparison correction ns: not significant; *: p ≤ 0.05; **: p < 0.01; ****: p < 0.0001

To investigate the localization of PICH on mitotic chromosomes treated with ICRF-193, mitotic cells were subjected to immunofluorescence staining. Synchronized cells were collected by mitotic shake off, treated with inhibitors for 20 minutes, then plated onto fibronectin coated coverslips. As seen in Western blot analysis, increased intensity of SUMO2/3 foci were enriched on the chromosomes, where they overlapped with TopoIIα foci upon ICRF-193 treatment (Figure 1B, enlarged images). Although, TopoIIα signal changed under Merbarone treatment, showing less punctate signal, no enrichment of SUMO2/3 foci were observed (Figure 1B). A novel observation showed that treatment with ICRF-193 caused a redistribution of PICH signal from all over the chromosome to an enrichment of foci on the chromosomes where they overlapped with SUMO2/3 foci (Figure 1C, enlarged images). Treatment with Merbarone did not affect PICH localization (Figure 1C). By outlining single chromosomes using the DNA signal in multiple images and then placing outlines on SUMO2/3 or TopoIIα channels the mean intensities of these signals were measured. Both TopoIIα and SUMO2/3 chromosome signal intensities were significantly higher in ICRF-193 treatment, but not in Merbarone treated cells (Figure 1D). PICH foci intensity was measured by using circles equal in size, the PICH foci intensity was found to be significantly increased in ICRF-193 treated cells (Figure 1D bottom graph). These data show that treatment with ICRF-193, but not Merbarone, causes increased TopoIIα SUMOylation and enrichment of PICH and SUMO2/3 foci on the chromosomes.

### SUMOylation is required for PICH enrichment in ICRF-193 treated cells

Although results obtained from inhibiting TopoIIα suggest that increased SUMOylation plays a critical role in PICH enrichment, the distinct effects of the different inhibitor treatments, for example differences in TopoII conformation, could also play a role. To determine if mitotic SUMOylation is critical for PICH enrichment in ICRF-193 treated cells we developed a novel method to inhibit mitotic SUMOylation in cells. First, we generated a fusion protein, called Py-S2, which consists of the N-terminal region of human PIASy, and the SENP2-catalytic domain (required for deSUMOylation) (Reverter and Lima, 2004; Ryu *et al.*, 2015; Sridharan *et al.*, 2015). The N-terminal region of PIASy localizes to mitotic chromosomes, in part, via its specific interaction with the RZZ (Rod-Zw10-Zwilch) complex (Ryu and Azuma, 2010). Thus, the fusion protein is expected to bring deSUMOylation activity where mitotic SUMOylation occurs on chromosomes by PIASy. As a negative control, we substituted a cysteine to alanine at position 548 of SENP2 (called Py-S2 Mut) to create a loss of function mutant (Reverter and Lima, 2004, 2006) (Figure 2A). The activity of the recombinant fusion proteins on chromosomal SUMOylation was verified in Xenopus egg extract (XEE) assays (Supplemental Figure S1). As predicted, addition of Py-S2 protein to XEE completely eliminated mitotic chromosomal SUMOylation. To our surprise, Py-S2 Mut protein stabilized SUMOylation of chromosomal proteins, thus acting as a dominant negative mutant against endogenous deSUMOylation enzymes. To express the fusion proteins in cells, we created inducible expression cell lines using the Tetracycline inducible system (Supplemental Figure S2) (Natsume *et al.*, 2016). We utilized CRISPR/Cas9 genome editing to integrate each of the fusion genes into the human H11 (hH11) safe harbor locus (Zhu *et al.*, 2014; Ruan *et al.*, 2015) in DLD-1 cells.

**Figure 2.**
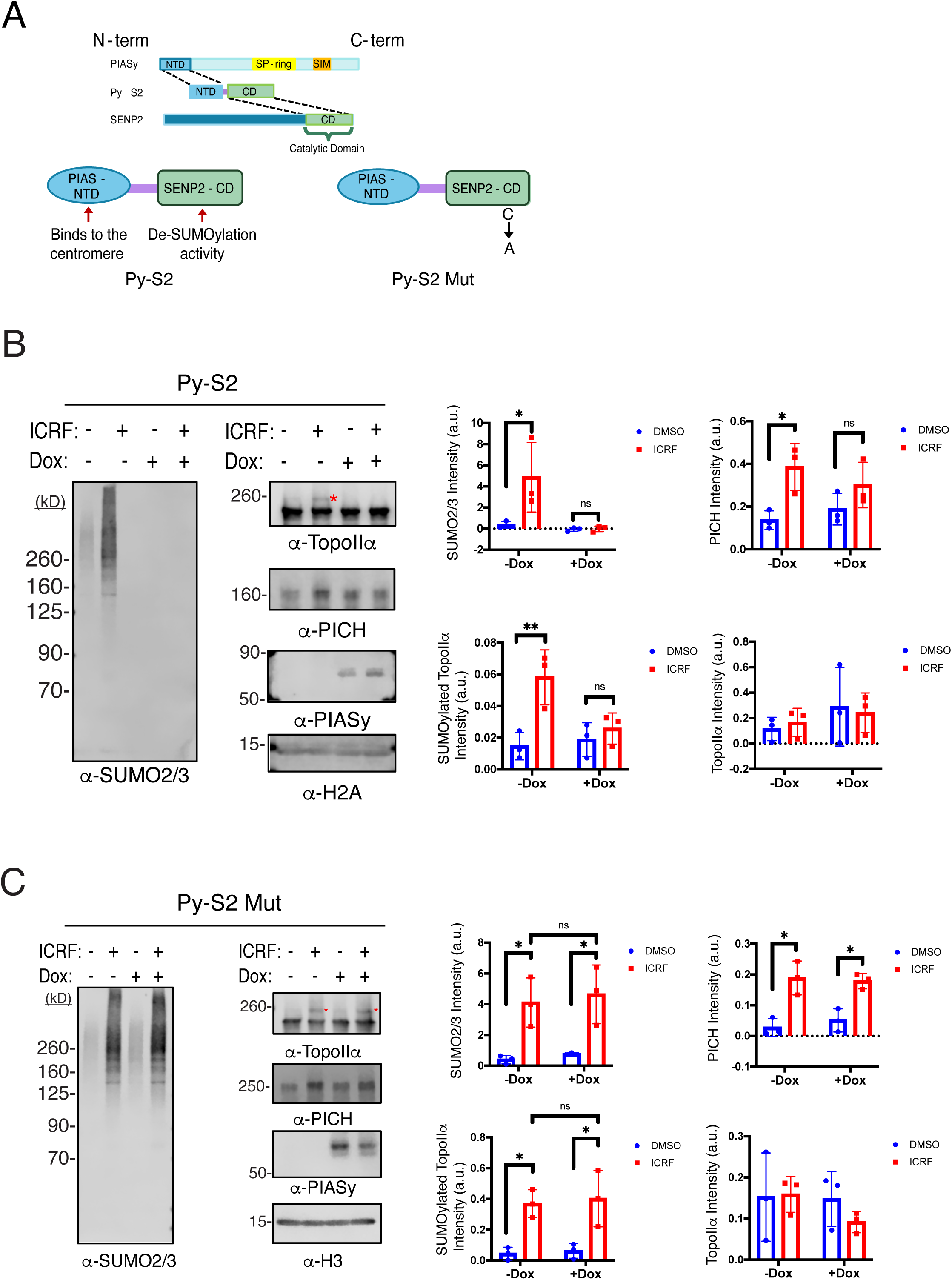
Modulating chromosomal SUMOylation affect chromosomal binding of PICH. **(A)** Schematic of fusion proteins generated for modulating SUMOylation on mitotic chromosomes. **(B)** Py-S2 expressing or non-expressing mitotic chromosomes were subjected to Western blotting with indicated antibodies. * indicates SUMOylated TopoIIα. Intensity of signals (a.u.) normalized by H2A are shown with mean value and standard deviation. p values for comparison from three experiments were calculated using a two-way ANOVA analysis of variance with Tukey multi-comparison correction; ns: not significant; *: p ≤ 0.05; **: p < 0.01 **(C)** Py-S2 Mut expressing or non-expressing mitotic chromosomes were isolated and subjected to Western blotting with indicated antibodies. * indicates SUMOylated TopoIIα. Intensity of signals (a.u.) normalized by H3 are shown with mean value with standard deviation. p values for comparison among three experiments were calculated using a two-way ANOVA analysis of variance with Tukey multi-comparison correction. ns: not significant; *: p ≤ 0.05

To test whether the Py-S2 fusion protein worked as expected, cells were synchronized, and doxycycline was added after release from a Thymidine block. After treatment with ICRF-193, chromosomes were isolated and subjected to Western blot analysis. The Py-S2 expressing cells had nearly undetectable levels of chromosomal SUMOylation as well as SUMOylated TopoIIα (Figure 2B). Intriguingly in Py-S2 expressing cells, PICH levels on chromosomes no longer showed a significant increase under ICRF-193 treatment, suggesting that the response of PICH to ICRF-193 depends on the cell’s ability to SUMOylate chromosomal proteins (Figure 2B, quantification). The role of SUMOylation in the enrichment of PICH on mitotic chromosomes is further supported by the Py-S2 Mut expressing cells. Western blot analysis of mitotic chromosomes expressing Py-S2 Mut revealed a slight increase of overall SUMOylation levels in the absence of ICRF-193 (Figure 2C lane 1 vs. lane 3, quantification). But, SUMOylated TopoIIα did not show a significant increase in either control or ICRF-193 treated conditions. This suggests that a similar stabilization of SUMOylation occurs in cells as was observed in the XEE assays, albeit with less penetrance. PICH levels were also slightly increased in the Py-S2 Mut expressing cells in the absence of ICRF-193 (Figure 2C lane 1 vs. lane 3, quantification).

To determine how deSUMOylation affects PICH and SUMO2/3 distribution on chromosomes, Py-S2 expressing mitotic cells were stained. Immunofluorescent analysis of Py-S2 expressing cells reiterated what was observed in Western blot analysis. By utilizing the same quantification methodology as Figure 1, measuring individual chromosome mean intensities, we saw that even under ICRF-193 treatment, Py-S2 expressing cells displayed significantly lower levels of SUMO2/3 signals on chromosomes (Figure 3A, quantification). PICH chromosomal intensity was also reduced in Py-S2 expressing cells, where no significant change occurred upon ICRF-193 treatment (Figure 3B, quantification). But, TopoIIα signals remained unaffected by inhibition of SUMOylation, agreeing with our previous observations in XEE assays (Azuma *et al.*, 2005) (Figure 3C, quantification). Next, the Py-S2 Mut expressing cells were synchronized and immunofluorescence analysis performed. The increase of both PICH and SUMO2/3 signals observed in Western blot quantification was mostly apparent with immunofluorescence analysis. We observed strong SUMO2/3 foci under ICRF-193 treatment in both Py-S2 Mut non-expressing and expressing cells. In control cells expressing Py-S2 Mut, quantification of SUMO2/3 mean intensity on chromosomes showed a significant increase and it was slightly higher than ICRF-193 treated Py-S2 expressing cells. This suggests that overall chromosomal SUMO2/3 modification is upregulated by Py-S2 Mut expression (Figure 4A, comparing black characters). The discrepancy between Western blot and quantification of immunofluorescence images imply that these increased SUMO2/3 modified proteins could be lost from chromosomal fractions during the isolation process for Western blotting. Intriguingly, in control cells expressing Py-S2 Mut we observed increased chromosomal PICH intensity as compared to ICRF-193 treated Py-S2 Mut expressing cells (Figure 4B, quantification). This mirrors the SUMO2/3 results and suggests that PICH interacts with SUMOylated chromosomal proteins in general and ICRF-193 treatment effects the formation of strong foci in both SUMO2/3 and PICH proteins. Similar to Figure 3C, TopoIIα localization and signal intensity did not change upon Py-S2 Mut expression (Figure 4C, quantification). In all, these data reinforce the indication that the enrichment of chromosomal PICH localization, including PICH foci formation in ICRF-193 treated cells, is dependent on increased SUMOylation.

**Figure 3.**
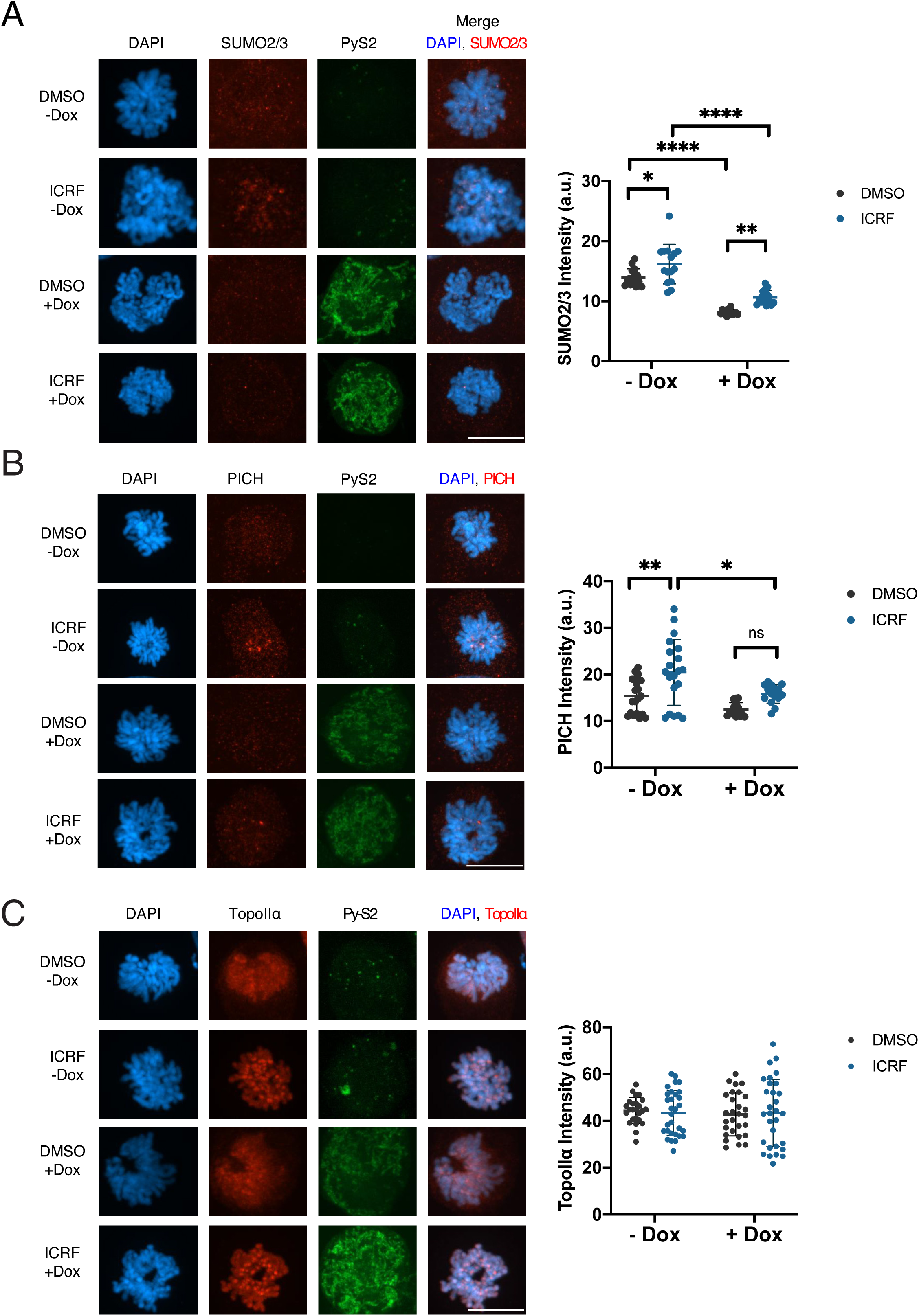
Decreased mitotic SUMOylation eliminates PICH response to ICRF-193. **(A)** Mitotic cells were fixed and stained with antibodies against: SUMO2/3 (red) and mNeon (green). DNA was stained by DAPI (blue). Scale bar = 11μm. The mean intensity of SUMO2/3 signals are shown. p values for comparison of all obtained values from three experiments were calculated using a two-way ANOVA analysis of variance with Tukey multi-comparison correction; *: p ≤ 0.05; **: p < 0.01; ****: p < 0.0001 **(B)** Mitotic cells were fixed and stained with antibodies against: PICH (red) and mNeon (green). DNA was stained by DAPI (blue). Scale bar = 11μm. The mean intensity of PICH signals are shown. p values for comparison of all obtained values from three experiments were calculated using a two-way ANOVA analysis of variance with Tukey multi-comparison correction; ns: not significant; *: p ≤ 0.05; **: p < 0.01 **(C)** Mitotic cells were fixed and stained with antibodies against: TopoIIα (red) and mNeon (green). DNA was stained by DAPI (blue). Scale bar = 11μm. The mean intensity of TopoIIα signals are shown. p values for comparison of all obtained values from three experiments were calculated using a two-way ANOVA analysis of variance with Tukey multi-comparison correction

**Figure 4.**
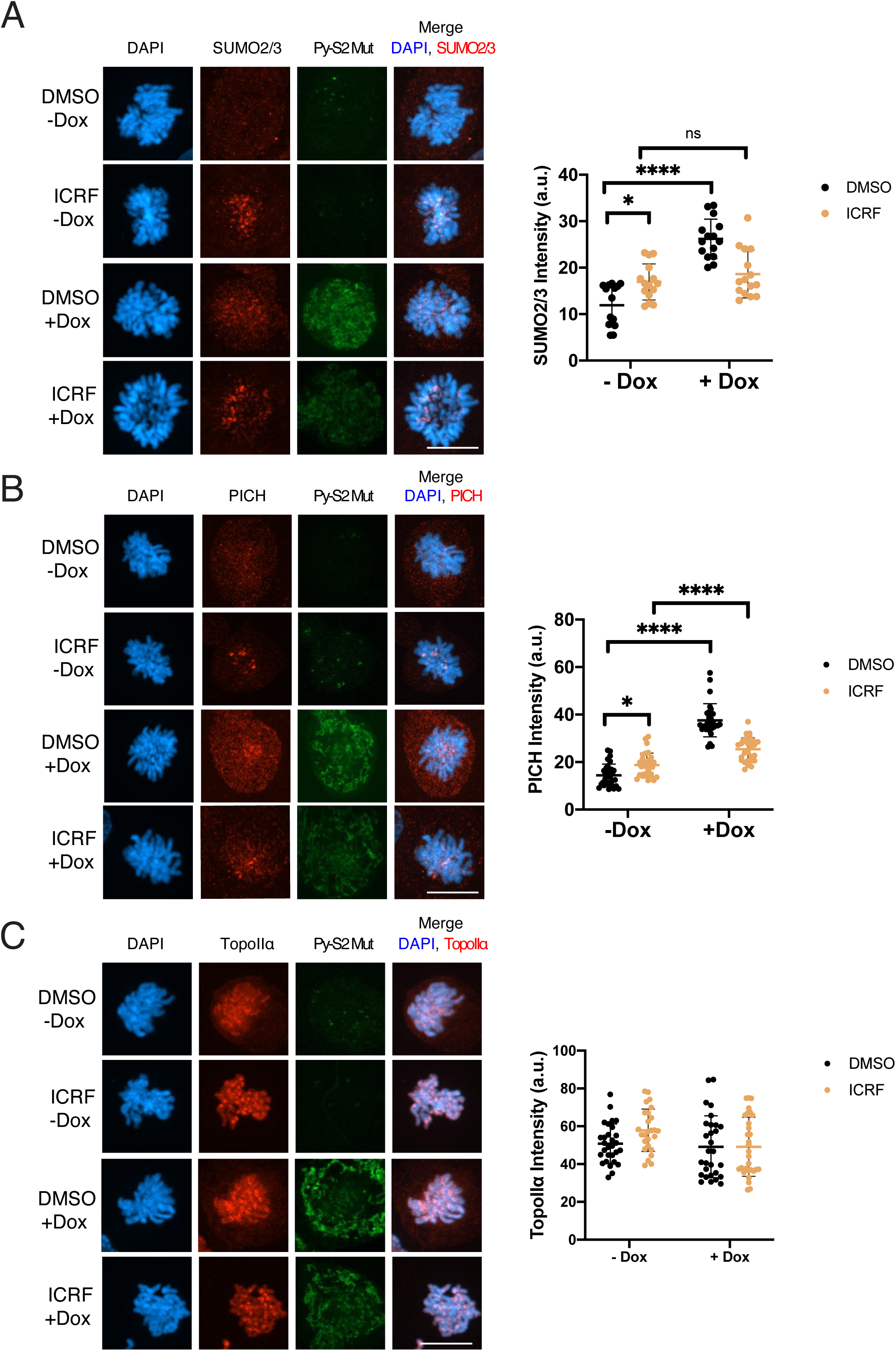
Mutant form of deSUMOylation enzyme promotes PICH and SUMO2/3 foci in both control and ICRF-193 treated cells. **(A)** Mitotic cells were fixed and stained with antibodies against: SUMO2/3 (red) and mNeon (green). DNA was stained by DAPI (blue). Scale bar = 11μm. The mean intensity of SUMO2/3 signals are shown. p values for comparison of all obtained values from three experiments were calculated using a two-way ANOVA analysis of variance with Tukey multi-comparison correction; ns: not significant; *: p ≤ 0.05; ****: p < 0.0001 **(B)** Mitotic cells were fixed and stained with antibodies against: PICH (red) and mNeon (green). DNA was stained by DAPI (blue). Scale bar = 11μm. The mean intensity of PICH signals are shown. p values for comparison of all obtained values from three experiments were calculated using a two-way ANOVA analysis of variance with Tukey multi-comparison correction; *: p ≤ 0.05; ****: p < 0.0001 **(C)** Mitotic cells were fixed and stained with antibodies against: TopoIIα (red) and mNeon (green). DNA was stained by DAPI (blue). Scale bar = 11μm. The mean intensity of TopoIIα signals are shown. p values for comparison of all obtained values from three experiments were calculated using a two-way ANOVA analysis of variance with Tukey multi-comparison correction

### Increased PICH levels observed in ICRF-193 treatment lost upon TopoIIα depletion

ICRF-193 treatment showed a distinct enrichment of SUMO2/3 and PICH foci unlike simply stabilizing SUMO2/3 modification on chromosomal proteins by Py-S2 Mut expression, therefore we tested whether the PICH response to ICRF-193 is due to TopoIIα SUMOylation. To accomplish this, we generated an mAID-TopoIIα cell line, which enables rapid and complete elimination of TopoIIα in the presence of auxin (Nishimura *et al.*, 2009; Natsume *et al.*, 2016). First, we established a cell line that has an integration of an auxin-dependent Ubiquitin E3 ligase, *OsTIR1* gene, at the promoter of a housekeeping (*RCC1)* gene (Supplemental Figure S3A-C) using CRISPR/Cas9 editing technology. The integration of the *OsTIR1* gene under the *RCC1* promoter achieved stable and low-level expression of the protein, thus minimized the non-specific degradation of AID-tagged proteins without auxin. Using the established OsTIR1 expressing DLD-1 cell line, DNA encoding a mAID-Flag tag was inserted into both TopoIIα loci (Supplemental Figure S4A-C). After 6-hour treatment with auxin, TopoIIα was degraded to undetectable levels in all cells analyzed (Supplemental Figure S4D and E). This rapid elimination allowed us to examine the effect of TopoIIα depletion in a single cell cycle.

To deplete TopoIIα, the cells were treated with auxin after release from a Thymidine block. After mitotic shake off and treatment with ICRF-193, isolated chromosomes were subjected to Western blotting with anti-SUMO2/3, anti-TopoIIα, and anti-PICH antibodies, and anti-H2A was used as a loading control. A slight overall increase of global SUMOylation was still observed in ΔTopoIIα cells treated with ICRF-193. Suggesting that ICRF-193 affects SUMOylation of other chromosomal proteins, such as TopoIIβ (Figure 5A). Notably, ΔTopoIIα cells treated with ICRF-193 showed no changes in PICH levels on the chromosomes. This suggests that increased levels of PICH seen in ICRF-193 treatment is a SUMOylated TopoIIα-dependent response (Figure 5A). Immunofluorescence analysis of PICH showed a significant reduction in signal on the chromosomes in ICRF-193 treatment as compared to cells with intact TopoIIα treated with ICRF-193. By measuring the mean PICH chromosomal signal intensity we saw a statistically significant decrease in mean PICH intensity in ICRF-193 treated ΔTopoIIα cells (Figure 5B, quantification). Similarly, ΔTopoIIα cells, showed a significant decrease in mean SUMO2/3 chromosome signal intensity under ICRF-193 treatment (Figure 5C, quantification). These results suggest that TopoIIα SUMOylation critically contributes to the enrichment of PICH on chromosomes under ICRF-193 treatment.

**Figure 5.**
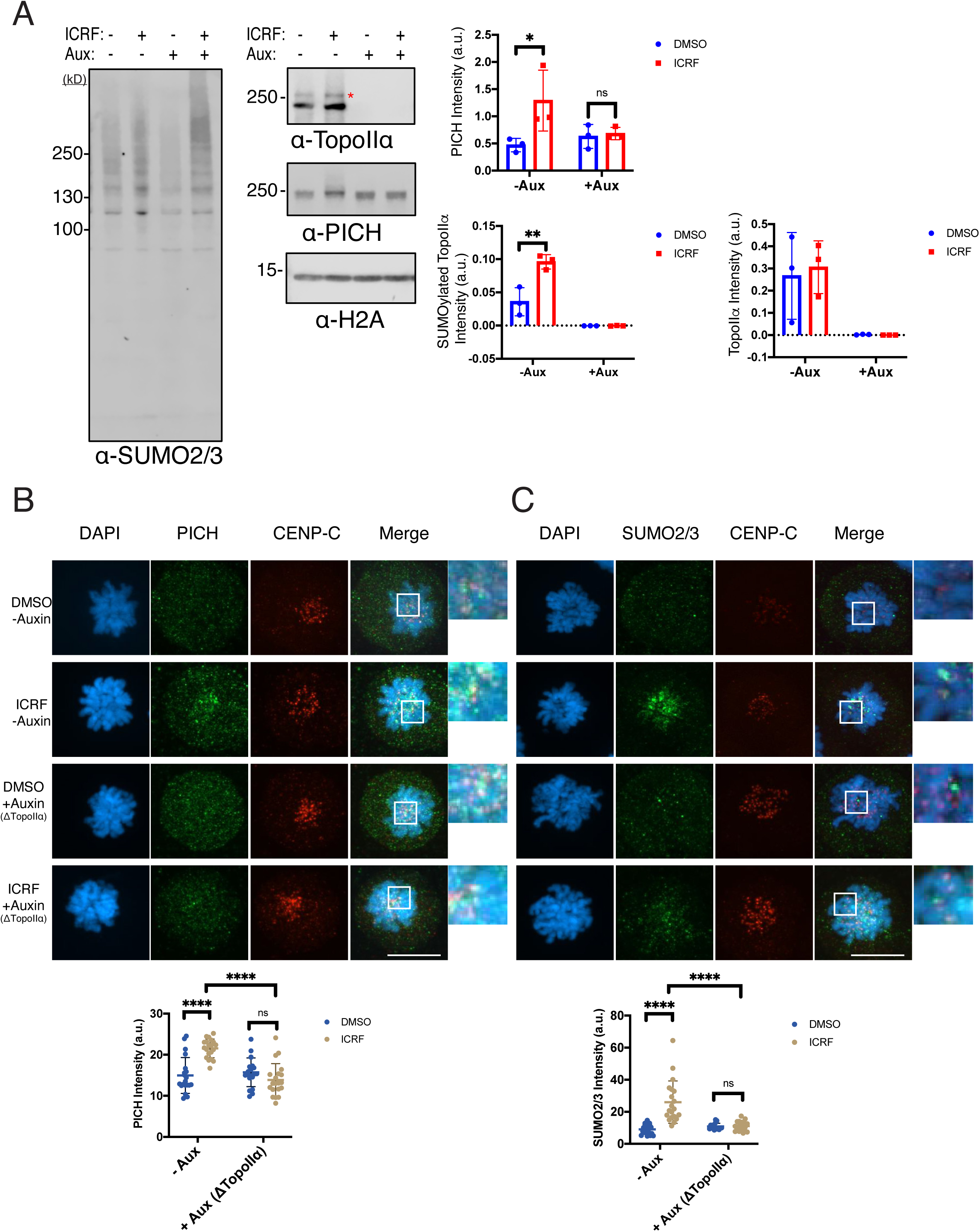
Depletion of TopoIIα attenuates SUMO2/3 modification and eliminates PICH response in ICRF-193 treated cells. **(A)** DLD-1 cells with endogenous TopoIIα tagged with a mAID were synchronized in mitosis and treated with DMSO (control) and ICRF-193. Auxin was added to the cells after release from Thymidine for 6 hours. Mitotic chromosomes were isolated and subjected to Western blotting with indicated antibodies. *indicates SUMOylated TopoIIα. Intensity of signals (a.u.) normalized by H2A are shown with mean value and standard deviation. p values for comparison from three experiments were calculated using a two-way ANOVA analysis of variance with Tukey multi-comparison correction; ns: not significant; *: p ≤ 0.05; **: p < 0.01 **(B)** Mitotic cells were fixed and stained with antibodies against: PICH (green), CENP-C (red). DNA was stained with DAPI (blue). Scale bar = 11μm. The white square indicates enlarged area. The mean intensity of PICH signals are shown. p values for comparison of all obtained values from three experiments were calculated using a two-way ANOVA analysis of variance with Tukey multi-comparison correction; ns: not significant; ****: p < 0.0001 **(C)** Mitotic cells were fixed and stained with antibodies against: SUMO2/3 (green), CENP-C (red). DNA was stained with DAPI (blue). Scale bar = 11μm. The white square indicates enlarged area. The mean intensity of SUMO2/3 signals are shown. p values for comparison of all obtained values from three experiments were calculated using a two-way ANOVA analysis of variance with Tukey multi-comparison correction; ns: not significant; ****: p < 0.0001

### Loss of PICH leads to enrichment of SUMOylated proteins

So far, the results indicate that PICH targets SUMOylated chromosomal proteins, in particular SUMOylated TopoIIα in ICRF-193 treated cells. Because the ability of PICH to interact with SUMO via its SUMO-interacting motifs is required for proper chromosome segregation, we wished to determine if PICH is required for regulating the association of SUMOylated proteins on chromosomes. To examine this, mAID-PICH cells were generated as described above for TopoIIα. After auxin was added to the cells for 6 hours, PICH levels became undetectable by Western blot and immunofluorescence analysis (Supplemental Figure S5A-E). To deplete PICH, auxin was added to the cells after release from a Thymidine block, then mitotic cells were collected by mitotic shake off. Isolated chromosomes were then subjected to Western blot analysis. Intriguingly, ΔPICH control cells showed a significant increase in SUMOylated TopoIIα compared to -Auxin cells, shown by the appearance of a second upshifted band marked by an asterisk (Figure 6A). Although not statistically significant, PICH-depletion showed an overall increase in SUMO2/3 signal in control cells by Western blot analysis (Figure 6A comparing lanes 1 and 3). Immunofluorescent staining further supported this novel role of PICH on SUMOylated chromosomal proteins. ΔPICH cells stained for SUMO2/3 showed a significant increase in mean chromosomal intensity as compared to control cells with intact PICH (Figure 6B, quantification). The discrepancy between Western blot and immunofluorescence image quantification suggests that these stabilized SUMO2/3 modified proteins on chromosomes under PICH-depletion are rather weakly associated to chromosomes and likely dissociate during isolation. In ΔPICH cells treated with ICRF-193 the mean intensity of SUMO2/3 was not significantly higher than cells with intact PICH. This suggests that PICH contributes to the attenuation or reorganization of SUMOylated proteins under normal conditions and ICRF-193 did not have an additive affect (Figure 6B, quantification). As compared to control cells with intact PICH, control ΔPICH cells showed a significant increase of mean TopoIIα chromosomal intensity (Figure 6C, quantification). This suggests that PICH can contribute to proper localization of TopoIIα on mitotic chromosomes regardless of ICRF-193. Together, the results suggest that PICH functions in the regulation and proper localization of SUMOylated chromosomal proteins.

**Figure 6.**
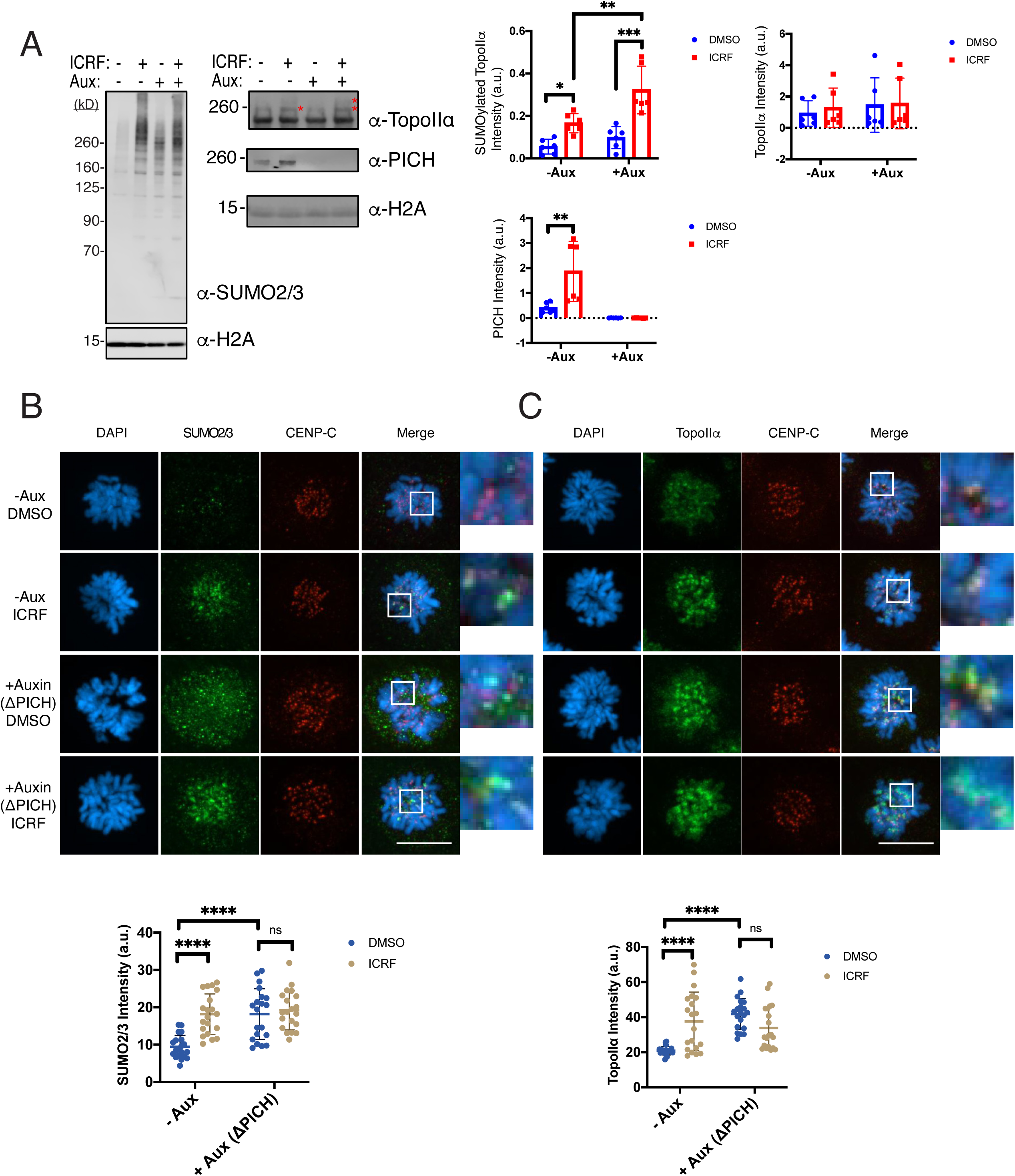
PICH-depleted chromosomes show increased levels of SUMOylated TopoIIα. **(A)** DLD-1 cells with endogenous PICH tagged with a mAID were synchronized in mitosis and treated with DMSO (control) and ICRF-193. Auxin was added to the cells after release from Thymidine for 6 hours. Mitotic chromosomes were isolated and subjected to Western blotting with indicated antibodies. *indicates SUMOylated TopoIIα. Intensity of signals (a.u.) normalized by H2A are shown with mean value and standard deviation. p values for comparison from three experiments were calculated using a two-way ANOVA analysis of variance and Tukey multi-comparison correction; ns: not significant; *: p ≤ 0.05; **: p < 0.01; ***: p < 0.001; ****: p < 0.0001. **(B)** Mitotic cells were fixed and stained with antibodies against: SUMO2/3 (green), CENP-C (red). DNA was stained with DAPI (blue). Scale bar = 11μm. The white square indicates enlarged area. The mean intensity of SUMO2/3 signals are shown. p values for comparison of all obtained values from three experiments were calculated using a two-way ANOVA analysis of variance and Tukey multi-comparison correction; ns: not significant; ****: p < 0.0001. **(C)** Mitotic cells were fixed and stained with antibodies against: TopoIIα (green), CENP-C (red). DNA was stained with DAPI (blue). Scale bar = 11μm. The white square indicates enlarged area. The mean intensity of TopoIIα signals are shown. p values for comparison of all obtained values from three experiments were calculated using a two-way ANOVA analysis of variance and Tukey multi-comparison correction; ns: not significant; ****: p < 0.0001.

### ATP-dependent translocase activity of PICH is required for regulating SUMOylated chromosomal proteins

To identify which function of PICH is required for the redistribution of SUMOylated proteins, we created a PICH-replacement cell line by combining mAID-mediated PICH depletion and inducible expression of exogenous PICH mutants. The mAID-PICH cells had CRISPR/Cas9 targeted integration of either Tet-inducible WT PICH-mCherry, an ATPase dead mutant (K128A-mCherry), or non-SUMO interacting form of PICH (d3SIM-mCherry) into the CCR5 safe harbor locus (Papapetrou and Schambach, 2016). After clonal isolation and validation (Supplemental Figure S6A-C), PICH-mCherry expression was tested in asynchronous cells by treating with auxin and doxycycline for 14 hours, and the whole cell lysates were used for Western blot analysis. Although the expression level of the exogenous proteins was variable, we were able to replace endogenous PICH with exogenous PICH (Figure 7A). We did observe variation of mCherry expression within each clonal isolate (Supplemental figure S6D) and this may explain the variation in expression levels observed in Western blot analysis. The PICH-replacement for mitotic cell analysis was achieved by incubating cells with auxin or auxin and doxycycline for 22 hours before mitotic shake off. The mitotic cells were treated with DMSO (control) and ICRF-193 then mitotic chromosomes were isolated. Western blot analysis was performed to determine how translocase activity and SIMs contribute to PICH binding to mitotic chromosomes (Figure 7B). The PICH WT-mCherry was observed to have a similar response to ICRF-193 as endogenous PICH, showing increased binding with ICRF-193 treatment. The K128A mutant also showed increased binding under ICRF-193 treatment. In contrast, the d3SIM mutant could not bind to chromosomes. This suggests that PICH SIMs are required for chromosomal association, which is consistent with our previous observations (Sridharan and Azuma, 2016).

**Figure 7.**
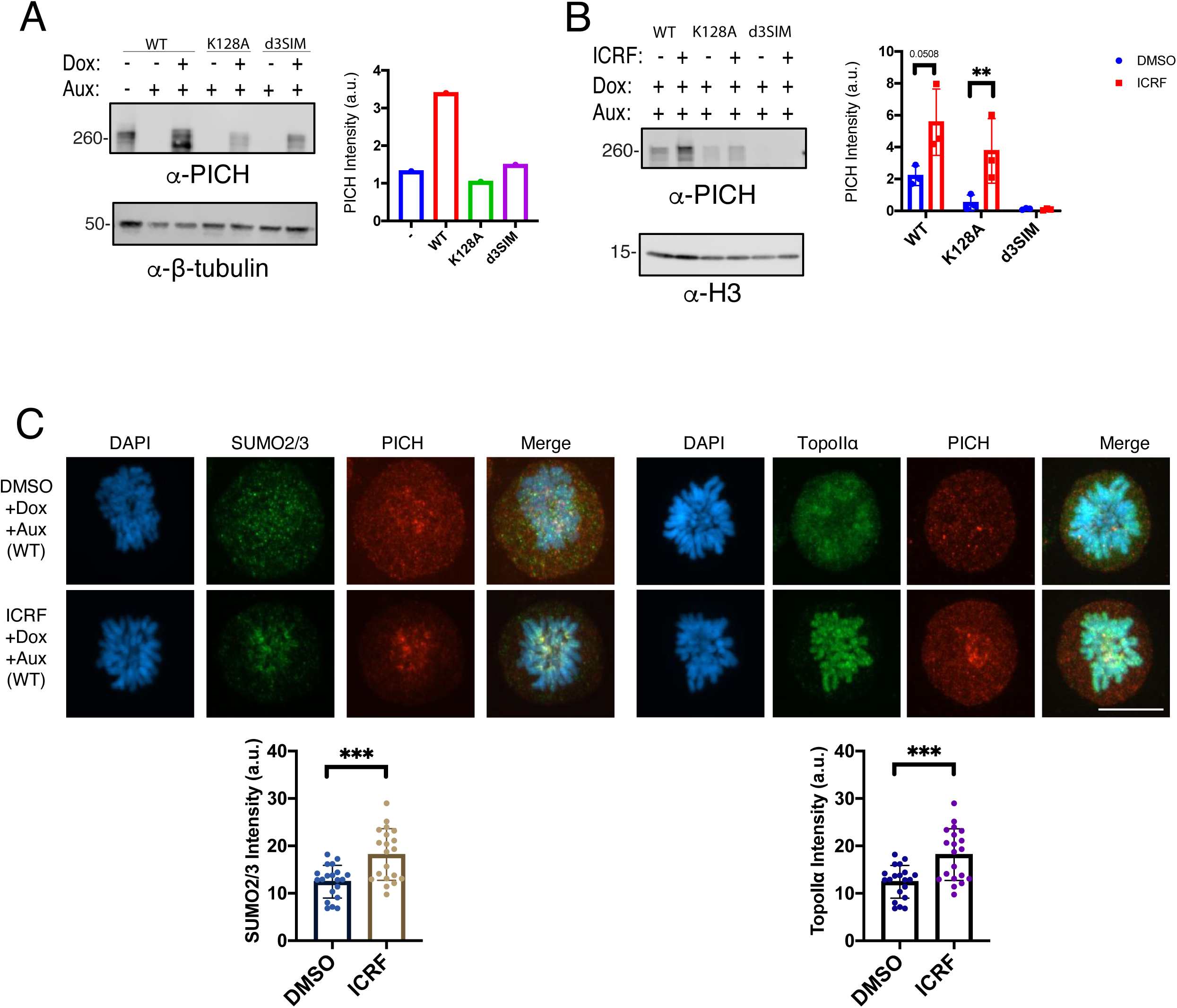
Expression of exogenous mCherry-tagged PICH functions similarly to endogenous PICH. **(A)** DLD-1 cells with endogenous PICH tagged with a mAID and exogenous PICH mCherry mutants were treated with auxin or auxin and doxycycline for 14 hours. Whole cell lysates were subjected to Western blotting with indicated antibodies. Intensity of PICH signals (a.u.) normalized by β-tubulin are shown. **(B)** DLD-1 cells with endogenous PICH tagged with a mAID and exogenous PICH mCherry mutants were treated with auxin or auxin and doxycycline for 22 hours. Mitotic chromosomes were isolated and subjected to Western blotting with indicated antibodies. Intensity of PICH signals (a.u.) normalized by H3 are shown with mean value and standard deviation. p values for comparison from three experiments were calculated using a Student t-test analysis of variance; ***: p < 0.001. **(C)** WT PICH mCherry mitotic cells were fixed and stained with antibodies against: SUMO2/3 (green), TopoIIα (green), mCherry (red). DNA was stained with DAPI (blue). Scale bar = 11μm. The mean intensity of SUMO2/3 or TopoIIα signals are shown. p values for comparison of all values from three experiments were calculated using a Student t-test analysis of variance; ***: p < 0.001.

To further examine how the PICH mutants affect chromosomal localization of SUMO2/3 and TopoIIα, immunofluorescent analysis of prometaphase cells was performed. PICH WT-mCherry showed the same staining patterns as endogenous PICH and its response to ICRF-193 was similar to Figure 1. Both SUMO2/3 and TopoIIα staining was consistent with that seen in Figure 1 (Figure 7C, quantification), further validating that mCherry tagged exogenous PICH functions the same as endogenous PICH.

When the K128A mutant, which cannot translocate DNA, was expressed strong mCherry foci were observed on the chromosomes regardless of ICRF-193 treatment. Importantly, these foci overlap with SUMO2/3 foci in all K128A cells observed, regardless of treatment (Figure 8A). This suggests that the PICH K128A mutant interacts with SUMOylated targets but due to its inability to translocate remains stably associated with the chromosomes where the SUMOylated proteins are located. SUMO2/3 mean intensity was observed to significantly increase in ICRF-193 treatment (Figure 8A, quantification). TopoIIα signals showed more diffused signal and foci formation was reduced in PICH-K128A expressing control cells. ICRF-193 treatment increased TopoIIα foci as in other conditions, however PICH foci did not show apparent colocalization with TopoIIα foci. Quantification of TopoIIα intensity on individual chromosomes showed no difference between control and ICRF-193 treatment (Figure 8A, quantification). This indicates that PICH translocase activity is required for the proper association and localization of TopoIIα on chromosomes. As observed by Western blot analysis, the PICH d3SIM-mCherry mutant did not show any chromosomal signal, but rather a diffuse signal was observed throughout the cell. Interestingly, even cells treated with ICRF-193 did not show an increased chromosomal SUMO2/3 signal, in turn there was no significant difference found in the mean SUMO2/3 chromosome signal intensity (Figure 8B, quantification). This was unexpected because depletion of PICH did not affect the increase of SUMO2/3 foci induced by ICRF-193 treatment. This observation suggests that the PICH d3SIM mutant has a negative effect on chromosomal SUMOylation, but the molecular mechanism of that phenomena is currently unidentified. TopoIIα in PICH d3SIM expressing cells was also affected showing an overall loss of chromosomal signal intensity where no significant difference was observed in ICRF-193 treatment (Figure 8B, quantification). This suggests that the SIM-dependent chromosomal association of PICH or reduced chromosomal SUMOylation is required for proper organization of mitotic chromosomes, which is required for proper distribution of TopoIIα on mitotic chromosomes.

**Figure 8.**
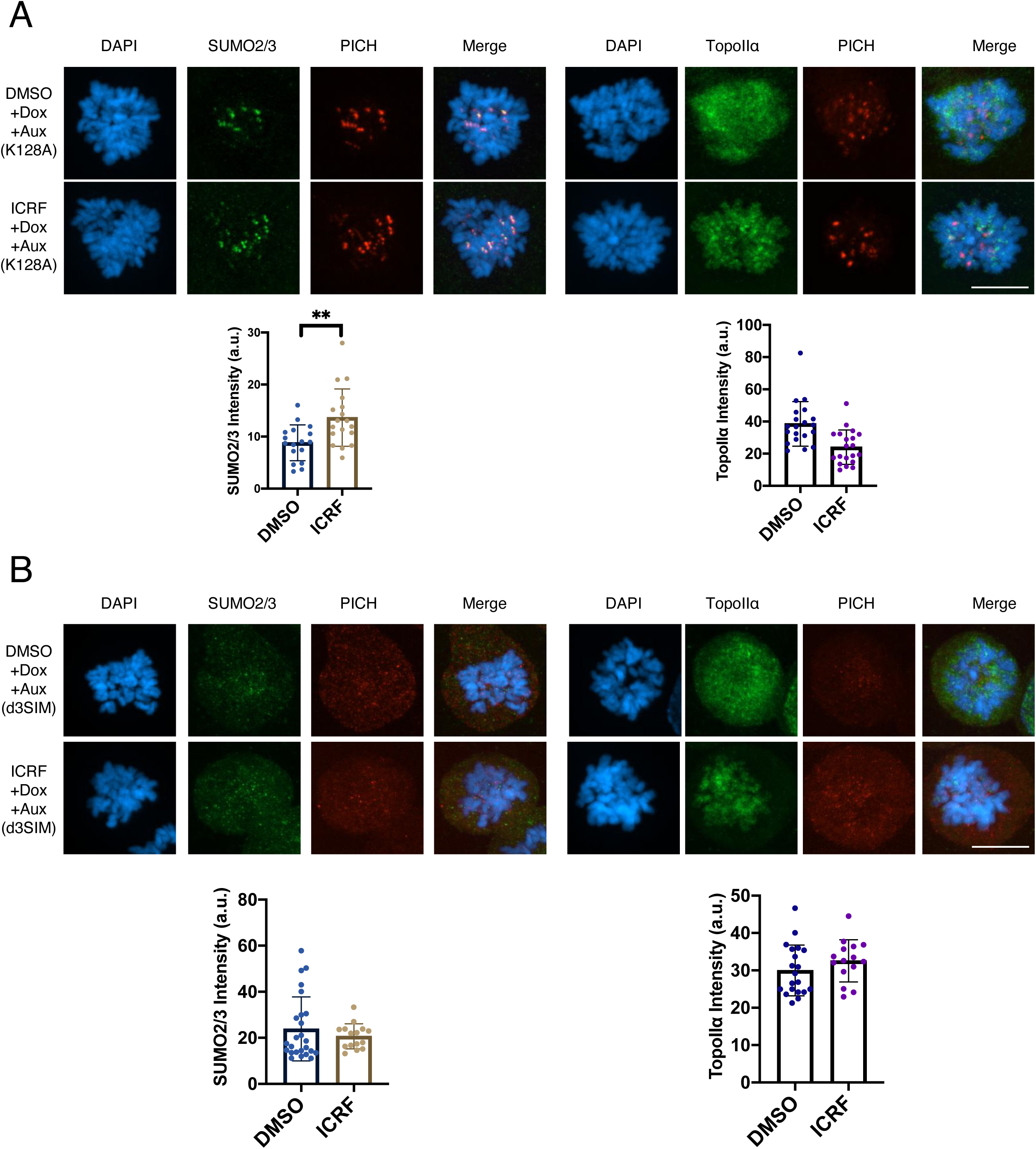
Translocase function of PICH is necessary for redistribution of SUMOylated proteins and SUMOylated TopoIIα on mitotic chromosomes. **(A)** K128A PICH mCherry mitotic cells were fixed and stained with antibodies against: SUMO2/3 (green), TopoIIα (green), mCherry (red). DNA was stained with DAPI (blue). Scale bar = 11μm. The mean intensity of SUMO2/3 signals are shown. p values for comparison of all values from three experiments were calculated using a Student t-test analysis of variance; **: p < 0.01. **(B)** d3SIM PICH mCherry mitotic cells were fixed and stained with antibodies against: SUMO2/3 (green), TopoIIα (green), mCherry (red). DNA was stained with DAPI (blue). Scale bar = 11μm. The mean intensity of TopoIIα signals are shown. p values for comparison of all values from three experiments were calculated using a Student t-test analysis of variance.

### PICH attenuates decatenation activity of SUMOylated TopoIIα through its SIMs

The cell-based assays suggest that PICH is required for proper organization of chromosomal SUMOylated proteins and SUMOylated TopoIIα is one of the targets under ICRF-193 treatment. To examine whether PICH can interact with SUMOylated TopoIIα and determine the potential role of their interaction, we performed an *in vitro* DNA decatenation assay to compare the effect of PICH on non-SUMOylated and SUMOylated TopoIIα (Figure 9A). Using the same conditions established in our previous study, recombinant *Xenopus laevis* TopoIIα was SUMOylated *in vitro*, then its DNA decatenation activity was analyzed by using catenated kDNA as the substrate (Ryu *et al.*, 2010b). The decatenation activity was measured by calculating the percentage of decatenated kDNA separated by gel electrophoresis. On average, 70% of kDNA is decatenated at the five and ten-minute time-point when non-SUMOylated TopoIIα is present in the reaction (Figure 9B PICH lanes marked by (i)). As we have previously shown, the decatenation activity of SUMOylated TopoIIα was reduced compared to non-SUMOylated TopoIIα (Figure 9B lanes marked by (ii)). Importantly, when we added PICH to each of the reaction at concentrations equimolar to TopoIIα (200nM), the decatenation activity of SUMOylated TopoIIα was further attenuated (Figure 9B marked by (iii), C). The reduction of decatenation activity of SUMOylated TopoIIα was statistically significant at both the five and ten-minute time-points (Figure 9C light grey bars). A dose-dependent effect of PICH on SUMOylated TopoIIα decatenation activity was observed but that was not the case for non-SUMOylated TopoIIα. The concentration of TopoIIα in the reaction was 200nM, and PICH significantly reduced decatenation activity of SUMOylated TopoIIα ranging between 200nM (equimolar) up to 400nM (Figure 9D, E). Only SUMOylated TopoIIα was inhibited by PICH dose-dependently which is distinct from the PICH/non-SUMOylated TopoIIα interaction.

**Figure 9.**
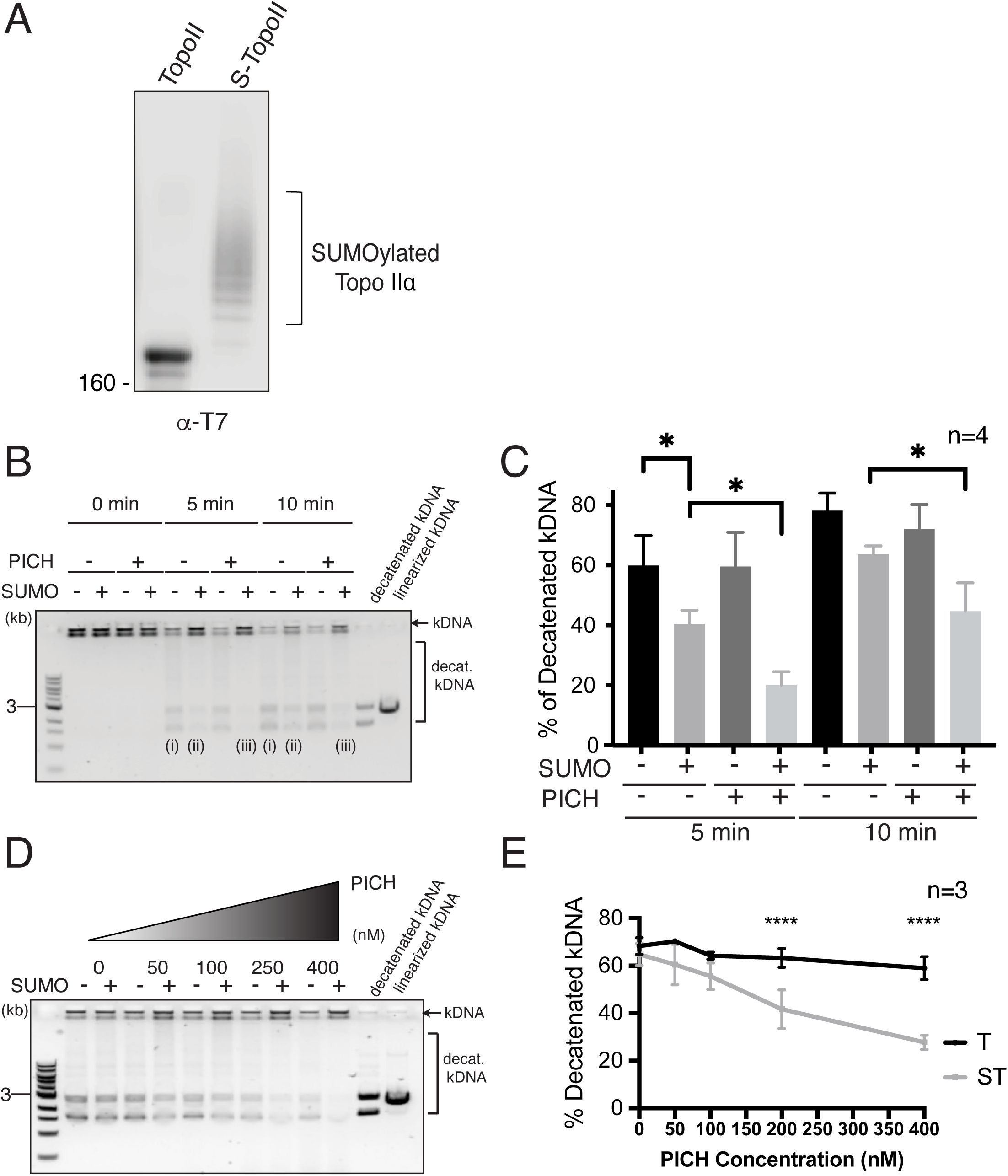
PICH inhibits SUMOylated TopoIIα decatenation activity. **(A)** Recombinant T7 tagged TopoIIα proteins were SUMOylated *in vitro*. Samples were subjected to Western blotting using anti-T7 tag antibody. The bracket indicates SUMOylated TopoIIα. **(B)** Representative gel after decatenation reactions with non-SUMOylated TopoIIα (**—** SUMO lane (i)) or SUMOylated TopoIIα (+ SUMO lane (ii)) (+PICH lane (iii)) Catenated kDNA is indicated by an arrow. The bracket indicates the decatenated kDNA species. **(C)** The decatenation activity of reactions in B was calculated as a percentage of decatenated kDNA. **(D)** Representative gel after decatenation reactions with SUMOylated and non-SUMOylated TopoIIα with increasing concentrations of PICH. Catenated kDNA is indicated by an arrow. The bracket indicates decatenated kDNA species. **(E)** The decatenation activity of SUMOylated (ST) and non-SUMOylated TopoIIα (T) in D was calculated as a percentage of decatenated kDNA. Statistical analysis of **C** (n=4) and **E** (n=3) were performed by using a two-way ANOVA analysis of variance with Tukey multi-comparison correction; p values for comparison among the experiments were calculated. ns: not significant; *: p ≤ 0.05; **: p < 0.01; ***: p < 0.001; ****: p < 0.0001

To determine which activity of PICH is required for inhibiting SUMOylated TopoIIα decatenation activity, we utilized a PICH mutant that has defects in either the SUMO-binding ability (PICH-d3SIM) or in translocase activity (PICH-K128A) (Figure 10A) (Sridharan and Azuma, 2016). If PICH/SUMO interaction is critical for inhibiting the decatenation activity of SUMOylated TopoIIα, the PICH-d3SIM mutant would lose its inhibitory function. In addition, we also expect that the PICH translocase activity deficient (PICH-K128A) mutant would lose its inhibitory function on SUMOylated TopoIIα, because this mutant could not remove SUMOylated TopoIIα from kDNA. Supporting our hypothesis, PICH-d3SIM lost its inhibitory function and SUMOylated TopoIIα decatenation activity returned to levels similar to no PICH addition (Figure 10C comparing ST to ST + PICH d3SIM). This suggests that direct SUMO/SIM interactions between PICH and SUMOylated TopoIIα play a key role in this inhibition. In contrast, the translocase deficient PICH mutant did not attenuate SUMOylated TopoIIα decatenation activity compared to WT PICH (Figure 10C comparing ST + PICH WT and ST + PICH K128A). Notably, neither of the PICH mutants showed any apparent effect on non-SUMOylated TopoIIα (Figure 10B) compared to PICH WT. This suggests that PICH binding to DNA does not inhibit the decatenation activity of TopoIIα, but rather it forms a complex with SUMOylated TopoIIα and prevents its decatenation activity. Taken together, our results suggest that PICH recognizes the SUMO moieties on TopoIIα through its SIMs to attenuate decatenation activity.

**Figure 10.**
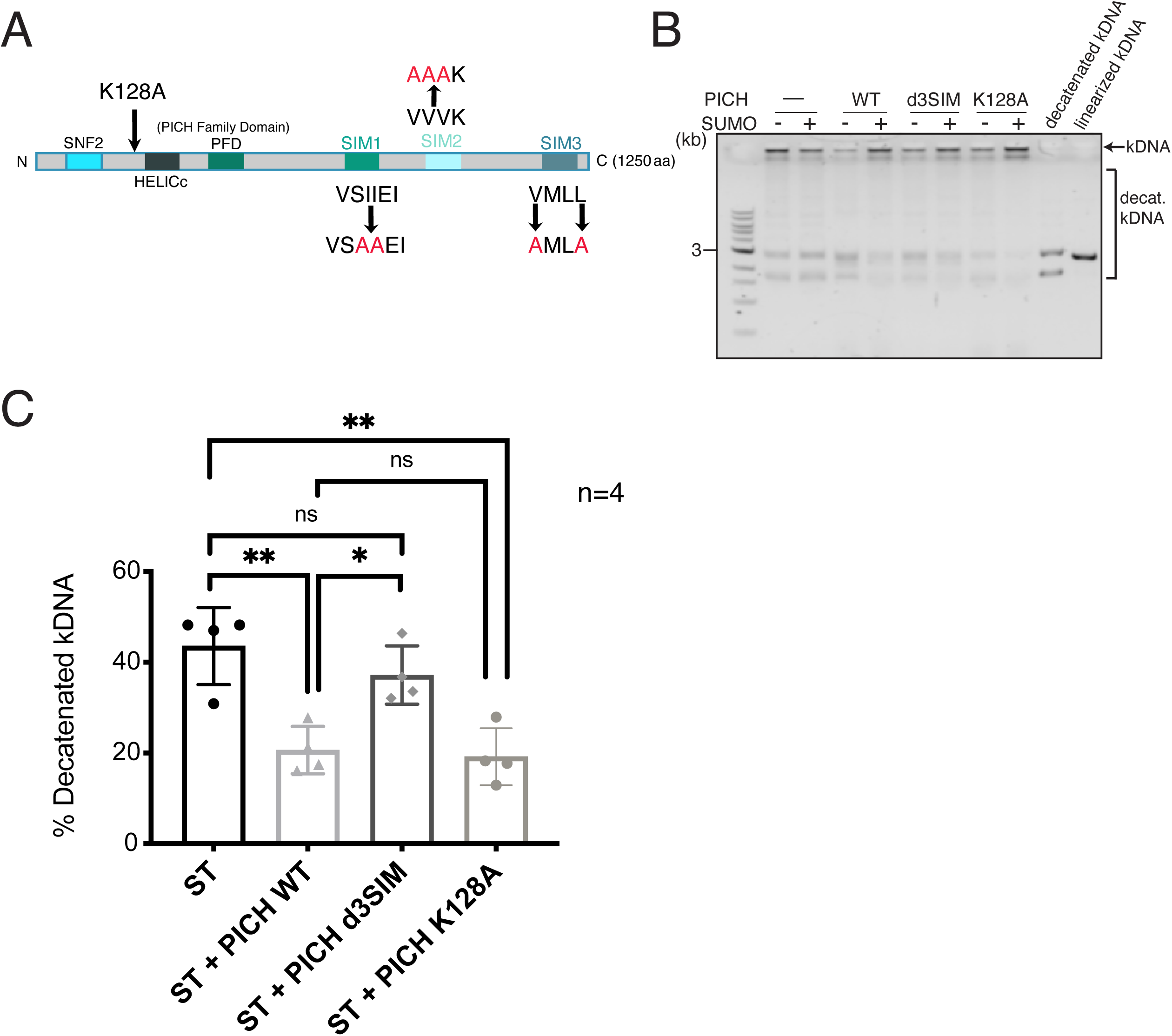
PICH SUMO-binding ability involved in suppression of SUMOylated TopoIIα decatenation activity. **(A)** Schematic of PICH protein with known functional motifs. The introduced mutations in SIMs and in the ATPase domain (K128A) are indicated. **(B)** Representative gel showing non-SUMOylated (-SUMO) and SUMOylated TopoIIα (+SUMO) activity with PICH WT, a non-SUMO-binding mutant (d3SIM), and a translocase deficient mutant (K128A) or no PICH protein (-PICH). Catenated kDNA is indicated with an arrow. The bracket indicates decatenated kDNA species. **(C)** Decatenation activity of SUMOylated TopoIIα (ST) with indicated PICH (ST: no PICH, ST + PICH WT: PICH wild-type, ST + PICH d3SIM: PICH-d3SIM mutant, and ST + PICH K128A: PICH-K128A mutant). Statistical analysis of **C** was performed by using a one-way ANOVA analysis of variance with Tukey multi-comparison correction; p values for comparison among four experiments were calculated. ns: not significant; *: p ≤ 0.05; **: p < 0.01.

In conclusion, our results show a novel function of PICH on the organization of SUMOylated chromosomal proteins during mitosis. This activity is dependent on PICH translocase activity and *in vitro* data suggests that SUMO interacting ability of PICH is important in the recognition of SUMOylated proteins (Figure 11).

**Figure 11.**
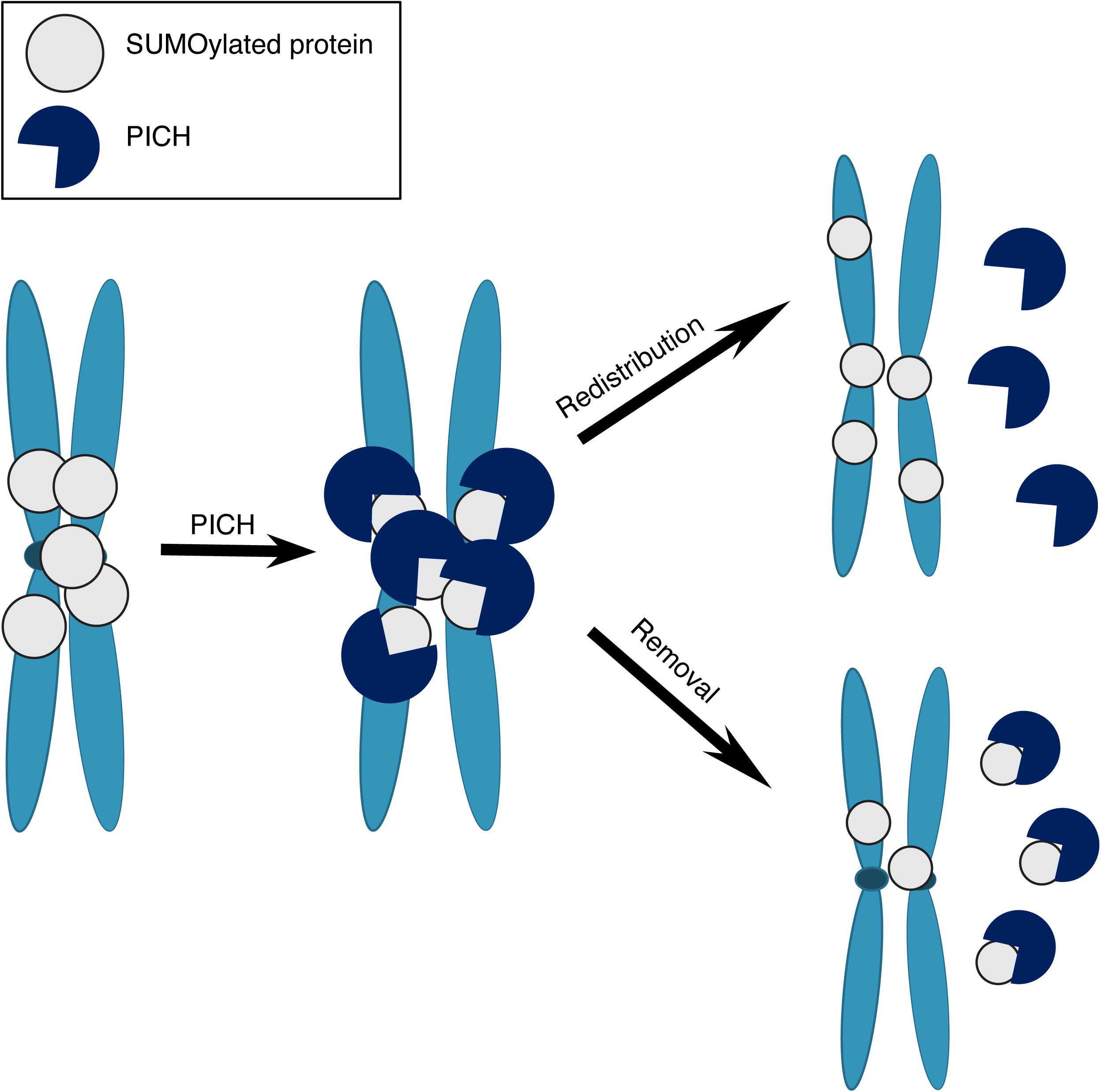
Model for demonstrating the role of PICH on regulating chromosomal SUMOylated proteins for proper chromosome segregation. SUMOylation plays a critical role in chromosome regulation and mitotic timing, this is due in part by regulating the activity and mediating binding of critical proteins. During mitosis proteins become SUMOylated and PICH recognizes and binds these proteins using its three SUMO interacting motifs, then using its translocase activity redistributes or removes SUMOylated proteins from the chromosomes and this enables proper chromosome segregation.

### Discussion

We previously demonstrated that both PICH DNA translocase activity and SUMO interacting ability are required for its essential function in proper chromosome segregation (Sridharan and Azuma, 2016). The results presented in this report provide the link between these two functions of PICH during mitosis. Collectively, the results indicate that PICH interacts with SUMOylated chromosomal proteins and increasing SUMOylation whether by modulating deSUMOylation enzyme or a specific inhibitor of TopoII mediates the enrichment of PICH foci on mitotic chromosomes. The PICH-replacement to mutant forms demonstrated that both DNA translocase activity and SUMO binding abilities are required for regulating proper localization of SUMOylated proteins on chromosomes.

### PICH targets and regulates chromosomal SUMOylated proteins using its SUMO binding ability and translocase activity

SUMOylation has been shown to play a role in complex assembly by mediating SUMO/SIM interactions (Lin *et al.*, 2006; Guzzo *et al.*, 2012; Pelisch *et al.*, 2017; Matmati *et al.*, 2018). It has been demonstrated that numerous proteins are SUMOylated on mitotic chromosomes (Schimmel *et al.*, 2014; Cubenas-Potts *et al.*, 2015; Huang *et al.*, 2016). Proper regulation of SUMOylation on chromosomal proteins is apparently key to promote faithful chromosome segregation shown by modulating enzymes for controlling SUMOylation (Hari *et al.*, 2001; Diaz-Martinez *et al.*, 2006; Cubenas-Potts *et al.*, 2013; Pelisch *et al.*, 2014). Our current study demonstrates that SUMOylated chromosomal proteins are targeted by PICH through its SIMs. Increased SUMO2/3 modification either by ICRF-193 (Figure 1) or expression of the deSUMOylation enzyme mutant (Figure 4) promotes enrichment of PICH and SUMO2/3 on chromosomes, and this suggests PICH efficiently targets SUMOylated chromosomal proteins including TopoIIα. Given the fact that PICH can interact with SUMO moieties (Sridharan *et al.*, 2015) using its three SIMs, this enrichment of PICH foci and SUMO2/3 foci suggests PICH can target multiple SUMOylated chromosomal proteins. More importantly, the translocase deficient mutant of PICH showed enrichment of SUMO2/3 foci on chromosomes without treatments to increase SUMOylation (Figure 8). Increased SUMO2/3 foci under expression of the mutant suggests that loss of translocase activity of PICH stabilized SUMOylated protein(s), presumably forming a stable complex on the chromosomes. Until now, the role of PICH DNA translocase activity in chromosome segregation has not been clearly determined on a cellular level. PICH primary structure suggests that it acts as a nucleosome remodeling enzyme, however, PICH has not been shown to have robust nucleosome remodeling activity towards nucleosomes composed of canonical histones (Ke *et al.*, 2011). Our observations imply that PICH could utilize its translocase activity for remodeling chromosomal SUMOylated proteins. Because the translocase deficient PICH mutant showed defective TopoIIα localization (shown in Figure 8), the organization of SUMO2/3 modified chromosomal proteins by PICH is required for proper formation of chromosomes. This is consistent with previous studies (Biebricher et al., 2013; Nielsen et al., 2015). One intriguing observation seen in the non-SUMO binding PICH mutant replaced cells is the loss of chromosomal SUMOylation and defects of TopoIIα localization. The SUMO binding deficient PICH mutant does not stably associate with mitotic chromosomes and compromises the response of increasing chromosomal SUMOylation under ICRF-193 treatment. This might indicate an unidentified function of non-chromosomal PICH on organization of mitotic chromosomes or reorganizing SUMO2/3 modified chromosomal proteins by PICH in the initial stages of mitosis is required for increased SUMO2/3 modification responding to ICRF-193. Identification of which SUMOylated chromosomal proteins are targeted by PICH and a more detailed analysis of chromosome organization in PICH-replaced cells will advance our understanding of the role of mitotic SUMOylation and the function of PICH in promoting faithful chromosome segregation.

### SUMOylated TopoIIα is a target of PICH under ICRF-193 treatment

Depleting TopoIIα abrogates the enrichment of PICH foci even in the presence of ICRF-193 (Figure 5), suggesting a specific role of PICH on SUMOylated TopoIIα in ICRF-193 treated cells. TopoIIα-depleted chromosomes also showed an ICRF-193 dependent increase in overall SUMOylation on chromosomes in Western blotting analysis, however, staining showed no clear increase of PICH foci. This suggests that PICH could more effectively target SUMOylated TopoIIα over other SUMOylated proteins under ICRF-193. It is notable that TopoIIα-depletion increases PICH binding with mitotic chromosomes even without upregulation of SUMOylation. This might represent the formation of PICH threads in TopoIIα-depleted prometaphase chromosomes (Antoniou-Kourounioti *et al.*, 2019), which are observed in ICRF-193 treated cells (Wang *et al.*, 2008). Therefore, increased PICH foci under ICRF-193 could be the result of the formation of PICH threads on prometaphase chromosomes. However, the results from Py-S2 expression (Figure 2 and 3) and PICH d3SIM mutant replacement (Figure 8) suggest that the increased PICH binding to chromosomes under ICRF-193 treatment is mainly controlled by the upregulation of SUMOylation. PICH binding with TopoIIα has been shown to increase TopoIIα activity *in vitro* (Nielsen *et al.*, 2015). In our assay, however, that increase was not clearly detected. This discrepancy might originate with the assay conditions, such as the existence of SUMO in the reaction. It is possible that excess SUMO in the reaction interacts with PICH, and that affects PICH interaction with unmodified TopoIIα. Once we determine how PICH interacts with unmodified TopoIIα, that will address the mechanism of activation of TopoIIα activity by PICH. In contrast, PICH binding to SUMOylated TopoIIα has different consequences, i.e. inhibition of decatenation activity (Figure 9). The inhibition of activity requires SIMs suggesting that direct interaction of PICH and SUMO moieties on TopoIIα is critical (Figure 10). The mechanism of how both WT PICH and translocase-deficient mutants similarly inhibit decatenation activity of SUMOylated TopoIIα is currently unclear. If we apply our model of “PICH as a SUMOylated protein remodeler” to this context, the mechanism of inhibition might be by removing SUMOylated TopoIIα from DNA. If that is the case, the translocase-deficient mutant could inhibit decatenation activity by forming a stable complex with SUMOylated TopoIIα on DNA, which can be predicted by the observation of stabilized SUMOylated protein on chromosomes in PICH-K128A replaced cells. Further analysis of the complex formation of PICH and SUMOylated TopoIIα *in vitro* or in cells is our next goal to elucidate the mechanism of this inhibition.

### Broader implications of the novel function of PICH as a SUMOylated protein remodeler

These novel findings lead to a more mechanistic understanding of the interaction between SUMOylated TopoIIα and PICH and provide insight into why PICH knockout cells were found to be sensitive to ICRF-193. PICH can increase TopoIIα decatenation activity *in vitro* and that helps to resolve tangled DNA during anaphase (Nielsen *et al.*, 2015). In addition, recent studies indicate that the translocase activity of PICH can be used to control the supercoiling status of DNA together with Topoisomerase IIIα (Bizard *et al.*, 2019). This increased supercoiling of DNA provides a more suitable substrate for TopoIIα and thus increases its decatenation activity. Both models can explain how PICH promotes decatenation on tangled DNA at centromeres to prevent UFB formation or resolve existing UFBs by stimulating TopoIIα activity. One unanswered question is how ICRF-193 mediated stalled TopoIIα is removed to prevent the formation of chromosome bridges. ICRF-193 treatment is known to induce a closed clamp conformation of TopoIIα with both detangled DNA strands bound within it (Roca *et al.*, 1994; Morris *et al.*, 2000). It is interesting to hypothesize from our current study that PICH SUMO-binding ability and translocase activity are able to recognize and bind SUMOylated TopoIIα and remove it from DNA. Analysis of PICH function using a TopoIIα-replaced cell line, utilizing the same methodology as the PICH mutant cell lines, will provide insight for this model. Recently, we demonstrated that ICRF-193 treatment resulted in a mitotic arrest in cells that requires SUMOylated TopoIIα and subsequent Aurora B activation (Pandey *et al.*, 2020). Because PICH can control SUMOylated TopoIIα on chromosomes, it is possible that PICH can control stalled TopoIIα-dependent mitotic checkpoint by attenuating SUMOylated TopoIIα on chromosomes. This can be tested using PICH depletion or replacement cell lines as well as modulating PICH activity in TopoIIα-replaced cell lines with a non-SUMOylatable mutant.

This novel role for PICH during mitosis leads to a better understanding of how chromosomal proteins are regulated by SUMOylation and how that might affect chromosome segregation when left unregulated. Although a precise molecular mechanism remains to be determined for the specific SUMOylated protein(s) targeted by PICH. One potential mechanism of how PICH could function with SUMOylated TopoIIα using both translocase activity and SUMO binding ability is presented from this study. A formal test of this model would greatly benefit the PICH field as its function during mitosis remains elusive. This would also shed light on how cells utilize PICH and TopoIIα to deal with the tangled DNA for proper chromosome segregation during mitosis.

## Materials and Methods

### Plasmids, constructs, and site-directed mutagenesis

The Py-S2 fusion DNA construct of human PIASy-NTD (amino acid 1-135) and SENP2-CD (amino acid 363-589) was created by fusion PCR method using a GA linker between the two fragments. Then, the Py-S2 fusion DNA fragment was subcloned into a recombinant expression pET28a plasmid at the BamHI/XhoI sites. To generate the Py-S2 Mut fusion DNA construct, substitution of Cysteine to Alanine at 548 in Py-S2 was introduced using a site-directed mutagenesis QuikChangeII kit (Agilent) by following the manufacturer’s instructions. hH11 locus and CCR5 locus targeting donor plasmids for inducible expression of Py-S2 proteins were created by modifying pMK243 (Tet-OsTIR1-PURO) plasmid (Natsume *et al.*, 2016). pMK243 (Tet-OsTIR1-PURO) was purchased from Addgene (#72835) and the OsTIR1 fragment was removed by BglII and MluI digestion, followed by an insertion of a multi-cloning site. Homology arms for each locus were amplified from DLD-1 genomic DNA using the primers listed in supplemental information. The Py-S2 fused with mNeon cDNA and PICH-mCherry fused cDNA were inserted at the MluI and SalI sites of the modified pMK243 plasmid. For CCR5 targeting plasmid, the antibiotics resistant gene was changed to Zeocin-resistant from Puromycin-resistant. The original plasmid for OsTIR1 targeting to RCC1 locus was created by inserting the TIR1 sequence amplified from pBABE TIR1-9Myc (Addgene #47328; (Holland *et al.*, 2012) plasmid, Blasticidin resistant gene (BSD) amplified from pQCXIB with ires-blast (Takara/Clontech), and miRFP670 amplified from pmiRFP670-N1 plasmid (Addgene #79987; (Shcherbakova *et al.*, 2016) into the pEGFP-N1 vector (Takara/Clontech) with homology arms for RCC1 C-terminal locus. Using genomic DNA obtained from DLD-1 cell as a template DNA, the homology arms were amplified using primers listed in supplemental information (Supporting information Table 1). Further, OsTIR1 targeting plasmid was modified by eliminating the miRFP670 sequence by PCR amplification of left homology arm and TIR/BSD/right homology arm for inserting into pMK292 obtained from Addgene (#72830) (Natsume *et al.*, 2016) using XmaI/BstBI sites. Three copies of codon optimized micro AID tag (50 amino-acid each (Morawska and Ulrich, 2013)) was synthesized by the IDT company, and hygromycin resistant gene/ P2A sequence was inserted upstream of the 3x micro AID sequence. The 3xFlag sequence from p3xFLAG-CMV-7.1 plasmid (Sigma) was inserted downstream of the AID sequence. The homology arms sequences for PICH N-terminal insertion and TopoIIα N-terminal insertion were amplified using primers listed in supplemental information (Table S1) from genomic DNA of DLD-1 cell, then inserted into the plasmid by using PciI/SalI and SpeI/NotI sites. In all of RCC1 locus, PICH locus, TopoIIα locus, CCR5 locus and hH11 locus genome editing cases, the guide RNA sequences listed in supplemental information (Table S1) were designed using CRISPR Design Tools from https://figshare.com/articles/CRISPR_Design_Tool/1117899 (Rafael Casellas laboratory, NIH) and http://crispr.mit.edu:8079 (Zhang laboratory, MIT) inserted into pX330 (Addgene #42230). Mutations were introduced in PAM sequences on the homology arms. The *X. laevis* TopoIIα cDNA and human PICH cDNA were subcloned into a pPIC 3.5K vector in which calmodulin-binding protein CBP-T7 tag sequences were inserted as previously described (Ryu *et al.*, 2010b; Sridharan and Azuma, 2016). All mutations in the plasmids were generated by site-directed mutagenesis using a QuikChangeII kit (Agilent) according to manufacturer’s instructions. All constructs were verified by DNA sequencing.

### Recombinant protein expression and purification, and preparation of antibodies

Recombinant TopoIIα and PICH proteins were prepared as previously described (Ryu *et al.*, 2010b; Sridharan and Azuma, 2016). In brief, the pPIC 3.5K plasmids carrying TopoIIα or PICH cDNA fused with Calmodulin binding protein-tag were transformed into the GS115 strain of *Pichia pastoris* yeast and expressed by following the manufacturer’s instructions (Thermo/Fisher). Yeast cells expressing recombinant proteins were frozen and ground with coffee grinder that contain dry ice, suspended with lysis buffer (50 mM Tris-HCl, pH 7.5, 150 mM NaCl, 2 mM CaCl_2_, 1 mM MgCl_2_, 0.1% Triton X-100, 5% glycerol, 1 mM DTT, complete EDTA-free Protease inhibitor tablet (Roche), and 10 mM PMSF). The lysed samples were centrifuged at 25,000 *g* for 40 min. To capture the CBP-tagged proteins, the supernatant was mixed with calmodulin-sepharose resin (GE Healthcare) for 90 min at 4°C. The resin was then washed with lysis buffer, and proteins were eluted with buffer containing 10 mM EGTA. In the case of PICH, the elution was concentrated by centrifugal concentrator (Amicon ultra with a 100kDa molecular weight cut-off). In the case of TopoIIα, the elution was further purified by Hi-trap Q anion-exchange chromatography (GE Healthcare). Recombinant Py-S2 proteins fused to hexa-histidine tag were expressed in Rossetta2 (DE3) (EMD Millipore/Novagen) and purified with hexa-histidine affinity resin (Talon beads from Takara/Clontech). Fractions by imidazole-elution were subjected to Hi-trap SP cation-exchange chromatography. The peak fractions were pooled then concentrated by centrifugal concentrator (Amicon ultra with a 30kDa molecular weight cut-off). The E1 complex (Aos1/Uba2 heterodimer), PIASy, Ubc9, dnUbc9, and SUMO paralogues were expressed in Rosetta2(DE3) and purified as described previously (Ryu *et al.*, 2010a).

To generate the antibody for human PICH, the 3’end (coding for amino acids 947∼1250) was amplified from PICH cDNA by PCR. The amplified fragment was subcloned into pET28a vector (EMD Millipore/Novagen) then the sequence was verified by DNA sequencing. The recombinant protein was expressed in Rossetta2(DE3) strain (EMD Millipore/Novagen). Expressed protein was found in inclusion body thus the proteins were solubilized by 8M urea containing buffer (20mM Hepes pH7.8, 300mM NaCl, 1mM MgCl2, 0.5mM TCEP). The solubilized protein was purified by Talon-resin (Clontech/Takara) using the hexa-histidine-tag fused at the N-terminus of the protein. The purified protein was separated by SDS-PAGE and protein was excised after Instant*Blue*™ (Sigma-Aldrich) staining. The gel slice was used as an antigen and immunization of rabbits was made by Pacific Immunology Inc., CA, USA. To generate the primary antibody for human TopoIIα, the 3’end of TopoIIα (coding for amino acids 1359∼1589) was amplified from TopoIIα cDNA by PCR. The amplified fragment was subcloned into pET28a and pGEX-4T vectors (GE Healthcare) then the sequence was verified by DNA sequencing. The recombinant protein was expressed in Rossetta2(DE3). The expressed protein was purified using hexa-histidine-tag and GST-tag by Talon-resin (Clontech/Takara) or Glutathione-sepharose (GE healthcare) following the manufacture’s protocol. The purified proteins were further separated by cation-exchange column. Purified hexa-histidine-tagged TopoIIα protein as used as an antigen and immunization of rabbits was made by Pacific Immunology Inc., CA, USA. For both PICH and TopoIIα antigens, antigen affinity columns were prepared by conjugating purified antigens (hexa-histidine-tagged PICH C-terminus fragment or GST-tagged TopoIIα C-terminus fragment) to the NHS-Sepharose resin following manufacture’s protocol (GE healthcare). The rabbit antisera were subjected to affinity purification using antigen affinity columns. Secondary antibodies used for this study and their dilution rates were: for Western blotting; Goat anti-Rabbit (IRDye^®^680RD, 1/20000, LI-COR) and Goat anti-Mouse (IRDye^®^800CW, 1/20000, LI-COR), and for immunofluorescence staining; Goat anti-mouse IgG Alexa Fluor 568 (#A11031, 1:500, Invitrogen), goat anti-rabbit IgG Alexa Fluor 568 (#A11036, 1:500, Thermo/Fisher), goat anti-rabbit IgG Alexa Fluor 488 (#A11034, 1:500, Thermo/Fisher), goat anti-guinea pig IgG Alexa Fluor 568 (#A21450, 1:500, Thermo/Fisher). Unless otherwise stated, all chemicals were obtained from Sigma-Aldrich.

### *In vitro* SUMOylation assays and decatenation assays

The SUMOylation reactions performed in the Reaction buffer (20 mM Hepes, pH 7.8, 100 mM NaCl, 5 mM MgCl_2_, 0.05% Tween 20, 5% glycerol, 2.5mM ATP, and 1 mM DTT) by adding 15 nM E1, 15 nM Ubc9, 45 nM PIASy, 500 nM T7-tagged TopoIIα, and 5 µM SUMO2-GG. For the non-SUMOylated TopoIIα control, 5 µM SUMO2-G mutant was used instead of SUMO2-GG. After the reaction with the incubation for one hour at 25°C, it was stopped with the addition of EDTA at a final concentration of 10mM. For the analysis of the SUMOylation profile of TopoIIα 3X SDS-PAGE sample buffer was added to reaction, and the samples were resolved on 8–16% Tris-HCl gradient gels (#XP08165BOX, Thermo/Fisher) by SDS-PAGE, then analyzed by Western blotting with HRP-conjugated anti-T7 monoclonal antibody (#T3699, EMD Millipore/Novagen).

Decatenation assays were performed in the Decatenation buffer (50 mM Tris-HCl, pH 8.0, 120 mM NaCl, 5 mM MgCl_2_, 0.5 mM DTT, 30 µg BSA/ml, and 2 mM ATP) with SUMOylated TopoIIα and non-SUMOylated TopoIIαn and with 6.2 ng/µl of kDNA (TopoGEN, Inc.). The resction was performed at 25°C with the conditions indicated in each of the figures. The reactions were stopped by adding one third volume of 6X DNA dye (30% glycerol, 0.1% SDS, 10 mM EDTA, and 0.2 µg/µl bromophenol blue). The samples were loaded on a 1% agarose gel containing SYBR™ Safe DNA Gel stain (#S33102, Invitrogen) with 1kb ladder (#N3232S, NEB), and electrophoresed at 100 V in TAE buffer (Tris-acetate-EDTA) until the marker dye reached the middle of the gel. The amount of kDNA remaining in the wells was measured using ImageStudio, and the percentage of decatenated DNA was calculated as (Intensity of initial kDNA [at 0 minutes incubation] - intensity of remaining catenated DNA)/Intensity of initial kDNA. Obtained percentages of catenated DNA was plotted and analyzed for the statistics by using GraphPad Prism 8 Software.

### Cell culture, Transfection, and Colony Isolation

Targeted insertion using the CRISPR/Cas9 system was used for all integration of exogenous sequences into the genome. DLD-1 cells were transfected with guide plasmids and donor plasmid using ViaFect™ (#E4981, Promega) on 3.5cm dishes. The cells were split and re-plated on 10cm dishes at ∼20% confluency, two days after, the cells were subjected to a selection process by maintaining in the medium in a presence of desired selection reagent (1μg/ml Blasticidin (#ant-bl, Invivogen), 400μg/ml Zeocin (#ant-zn, Invivogen), 200μg/ml Hygromycin B Gold (#ant-hg, Invivogen)). The cells were cultured for 10 to 14 days with a selection medium, the colonies were isolated and grown in 48 well plates, and prepared Western blotting and genomic DNA samples to verify the insertion of the transgene. Specifically, for the Western blotting analysis, the cells were pelleted, 1X SDS PAGE sample buffer was added, and boiled/vortexed. Samples were separated on an 8-16% gel and then blocked with Casein and probed using the indicated antibody described in each figure legend. Signals were acquired using the LI-COR Odyssey Fc imager. To perform genomic PCR, the cells were pelleted, genomic DNA was extracted using lysis buffer (100mM Tris-HCl pH 8.0, 200mM NaCl, 5mM EDTA, 1% SDS, and 0.6mg/mL proteinase K (#P8107S, NEB)), and purified by ethanol precipitation followed by resuspension with TE buffer containing 50ug/mL RNase A (#EN0531,ThermoFisher). Primers used for confirming the proper integrations are listed in the supplemental information.

To establish AID cell lines, as an initial step, the *Oryza sativa* E3 ligase (OsTIR1) gene was inserted into the 3’ end of a housekeeping gene, RCC1, using CRISPR/Cas9 system in the DLD-1 cell line. The RCC1 locus was an appropriate locus to accomplish the modest but sufficient expression level of the OsTIR1 protein so that it would not induce a non-specific degradation without the addition of Auxin (Supplemental Figure S3). We then introduced DNA encoding for AID-3xFlag tag into the TopoIIα or PICH locus using CRISPR/Cas9 editing into the OsTIR1 expressing parental line (Supplemental Figure S4 and S5). The isolated candidate clones were subjected to genomic PCR and Western blotting analysis to validate integration of the transgene. Once clones were established and the transgene integration was validated, the depletion of the protein in the auxin-treated cells was confirmed by Western blotting and immunostaining.

Introducing DNA encoding Tet inducible PICH mCherry into the CCR5 locus or inducible Py-S2 into hH11 were made by CRISPR/Cas9 editing into the desired locus (Supplemental Figure S2 and S6). The OsTIR1 expressing, mAID PICH parental cell line was used for introduction of the PICH mCherry mutants targeted to the CCR5 locus. The isolated candidate clones were subjected to genomic PCR and Western blotting analysis to validate integration of the transgene. Once clones were established and the transgene integration was validated, the expression of the transgenes was confirmed by the addition of doxycycline.

### Xenopus egg extract assay for mitotic chromosomal SUMOylation analysis

Low speed cytostatic factor (CSF) arrested Xenopus egg extracts (XEEs) and demembraned sperm nuclei were prepared following standard protocols (Murray, 1991; Powers *et al.*, 2001). To prepare the mitotic replicated chromosome, CSF extracts were driven into interphase by adding 0.6mM CaCl_2_. Demembraned sperm nuclei were added to interphase extract at 4000 sperm nuclei/μl, then incubated for ∼60 min to complete DNA replication confirmed by the morphology of nuclei. Then, equal volume of CSF XEE was added to the reactions to induce mitosis. To confirm the activities of Py-S2 proteins on mitotic SUMOylation, the Py-S2 proteins or dnUbC9 were added to XEEs at a final concentration of 30nM and 5μM, respectively, at the onset of mitosis-induction. After mitotic chromosome formation was confirmed by microscopic analysis of condensed mitotic chromosomes, chromosomes were isolated by centrifugation using 40% glycerol cushion as previously described (Yoshida *et al.*, 2016) then the isolated mitotic chromosomes were boiled in SDS-PAGE sample buffer. Samples were resolved on 8-16% gradient gels and subjected to Western blotting with indicated antibodies. Signals were acquired using LI-COR Odyssey Fc digital imager and the quantification was performed using Image Studio Lite software. The following primary antibodies were used for Western blotting: Rabbit anti-Xenopus TopoIIα (1:10,000), Rabbit anti-Xenopus PARP1 (1:10,000), Rabbit anti-SUMO2/3 (1:1,000) (all prepared as described previously (Ryu *et al.*, 2010a)), anti-Histone H3 (#14269, Cell Signaling).

### Preparation of mitotic cells and chromosome isolation

DLD-1 cells were grown in McCoy’s 5A 1x L-glutamine 10% FBS media for no more than 10 passages. To analyze mitotic chromosomes, cells were synchronized by Thymidine/Nocodazole cell cycle arrest protocol. In brief, cells were arrested with 2mM Thymidine for 17 hours, were released from the Thymidine block by performing three washes with non-FBS containing McCoy’s 5A 1x L-glutamine media and placed in fresh 10%FBS containing media. 6 hours after the Thymidine release, 0.1ug/mL Nocodazole was added to the cells for 4 additional hours, mitotic cells were isolated by performing a mitotic shake-off and washed 3 times using McCoy’s non-FBS containing media to release from Nocodazole. The cells were then resuspended with 10% FBS containing fresh media and 7uM of ICRF-193, 40uM Merbarone, or equal volume DMSO, were plated on Fibronectin coated cover slips, and incubated for 20 minutes (NEUVITRO, #GG-12-1.5-Fibronectin). To isolate mitotic chromosomes, the cells were lysed with lysis buffer (250mM Sucrose, 20mM HEPES, 100mM NaCl, 1.5mM MgCl_2_, 1mM EDTA, 1mM EGTA, 0.2% TritonX-100, 1:2000 LPC (Leupeptin, Pepstatin, Chymostatin, 20mg each/ml in DMSO; Sigma-Aldrich), and 20mM Iodoacetamide (Sigma-Aldrich #I1149)) incubated for 5 minutes on ice. Lysed cells were then placed on a 40% glycerol containing 0.25% Triton-X-100 cushion, and spun at 10,000xg for 5 minutes, twice. Isolated chromosomes were then boiled with SDS-PAGE sample buffer, resolved on an 8-16% gradient gel and subjected to Western blotting with indicated antibodies. Signals of the blotting were acquired using the LI-COR Odyssey Fc machine.

The following primary antibodies were used for Western blotting: Rabbit anti-PICH (1:1,000), Rabbit anti-TopoIIα (1:20,000) (both are prepared as described above), Rabbit anti-SUMO2/3 (1:1,000), Rabbit anti-Histone H2A (1:2,000) (#18255, Abcam), Rabbit anti-Histone H3 (1:2,000) (#14269, Cell Signaling), Rabbit anti-PIASy (1:500) (as described in (Azuma *et al.*, 2005)), Mouse anti-β-actin (1:2,000) (#A2228, Sigma-Aldrich), Mouse anti-myc (1:1,000) (#9E10, Santa Cruz), Mouse anti-β-tubulin (1:2,000) (#, Sigma-Aldrich), Mouse anti-Flag (1:1,000) (#F1804, Sigma-Aldrich).

### Cell fixation and staining

To fix the mitotic cells on fibronectin coated cover slips, cells were incubated with 4% paraformaldehyde for 10 minutes at room temperature, and subsequently washed three times with 1X PBS containing 10mM Tris-HCl to quench PFA. Following the fixation, the cells were permeabilized using 100% ice cold Methanol in -20°C freezer for 5 minutes. Cells were then blocked using 2.5% hydrolyzed gelatin for 30 minutes at room temperature. Following blocking the cells were stained with primary antibodies for 1 hour at room temperature, washed 3 times with 1X PBS containing 0.1% tween20, and incubated with secondary for 1 hour at room temperature. Following secondary incubation cells were washed 3 times with 1x PBS-T and mounted onto slide glass using VECTASHIELD^®^ Antifade Mounting Medium with DAPI (#H-1200, Vector laboratory) and sealed with nail polish. Images were acquired using an UltraView VoX spinning disk confocal system (PerkinElmer) mounted on an Olympus IX71 inverted microscope. It was equipped with a software-controlled piezoelectric stage for rapid Z-axis movement. Images were collected using a 60 × 1.42 NA planapochromatic objective (Olympus) and an ORCA ERAG camera (Hamamatsu Photonics). Solid state 405, 488, and 561 nm lasers were used for excitation. Fluorochrome-specific emission filters were used to prevent emission bleed through between fluorochromes. This system was controlled by Volocity software (PerkinElmer). Minimum and maximum intensity cutoffs (black and white levels) for each channel were chosen in Volocity before images were exported. Images are presented as extended focus. No other adjustments were made to the images. The cell mages of supplemental figures were acquired using the Plan Apo 100x/1.4 objective lens on a Nikon Ti Eclipse microscope equipped Exi Aqua CCD camera (Q imaging) or a Nikon TE2000-U equipped PRIME-BSI CMOS camera (Photometrics) with MetaMorph imaging software. Figures were prepared from exported images in Adobe illustrator.

The following primary antibodies were used for staining: Rabbit anti-PICH 1:800, Rabbit anti-human TopoIIα 1:1000 (both are prepared as described above), Mouse anti-human TopoIIα 1:300 (#Ab 189342, Abcam), Mouse anti-SUMO2/3 (#12F3, Cytoskeleton Inc), Guinea Pig anti-SUMO2/3 (1:300) (prepared as previously described (Ryu et al., 2010), and Rat anti-RFP (#RMA5F8, Bulldog Bio Inc).

### Statistical analysis of immunofluorescent images

Quantification was performed by measuring at least 5 chromosomes per treatment across three individual experiments. This was done by outlining the chromosome and superimposing that drawing onto the other channels before measuring. All data provided are mean intensities. PICH foci intensity were measured in Figure 1 by creating a 10×10 circle on chromosome ends and superimposing circles onto PICH channels and measuring mean intensities. At least 5 chromosome ends were measured per treatment across three individual experiments.

### Statistical analysis

All statistical analyses were performed with either 1- or 2-way ANOVA, followed by the appropriate post-hoc analyses using GraphPad Prism 8 software. Graphs are presented as mean with standard deviation.

### Animal use

For XEE assay, frog eggs were collected from a mature female *Xenopus laevis*, and sperm was obtained from matured male *Xenopus laevis*. The animal use protocol for the *Xenopus laevis* studies was approved by University of Kansas IACUC.

## Abbreviations

TopoIIα: Topoisomerase IIα
PICH: Polo-like kinase interacting checkpoint helicase
SPR: Strand passage reaction
SUMO: Small ubiquitin-like modifier
XEE: Xenopus egg extract
CSF: Cytostatic factor
dnUbc9: dominant negative E2 SUMO-conjugating enzyme
SENP: Sentrin-specific protease
PIAS: Protein inhibitor of activated STAT
SIM: SUMO-interacting-motif

## Acknowledgements

We thank Drs. M. Azuma, V. Paolillo and B. R. Oakley at the University of Kansas for the use of their microscopes and for technical assistance during microscope and software usage. We also thank Dr. D. Clarke at the University Minnesota and Dr. Y. Yamashita at the University of Michigan for the critical reading of the manuscript and comments on this project. This work was supported by NIH/NIGMS, GM112893 and, in part, by KUCC/CB pilot grant (KAN1000623). The establishment of AID-mediated knockdown system was supported V. Aksenova, A. Arnaoutov and M. Dasso whom are supported by the National Institute for Child Health and Human Development Intramural projects Z01 HD008954 and ZIA HD001902.

## Author Contributions

VH conducted almost all of the experiments, created the AID fused PICH cell line and PICH-replaced cell lines, prepared figures, and drafted the manuscript. HP prepared DNA constructs for genome editing, created CRISPR/Cas9 genome edited for inducible expression of de-SUMOylation enzyme and for Os-TIR1 expressing DLD-1 cell line, created AID fused TopoIIα cell lines, and performed XEE assay for validation of Py-S2 proteins. NP conducted experiments for initial validation of the genome edited cell lines expressing Py-S2. BL performed initial analysis of immunofluorescent images in Figure 5. VA, AA, and MD established AID-mediated degradation system by optimizing Os-TIR1 integration locus and creating constructs for genome editing by CRISPR/Cas9 for that system. YA designed the study, supervised project, and wrote the manuscript.

## Figure legend

**Supplemental Figure S1. Testing SUMO modulating proteins in the *Xenopus laevis* egg extract system.**

**(A)** Recombinant Py-S2 or Py-S2 Mut proteins were added to *Xenopus laevis* egg extract upon induction of mitosis, and the chromosomes were isolated. Chromosome samples were subjected to Western blotting with anti-SUMO2/3 antibody.

**(B)** Chromosome samples in A were subjected to Western blotting with anti-Xenopus TopoIIα antibody to detect both TopoIIα (∼160kDa) and SUMOylated TopoIIα (marked with red asterisks), and anti-Xenopus PARP1 antibody to detect both PARP1 (∼100kDa) and SUMOylated PARP1 (marked with red asterisks). Anti-histone H3 antibody was used as a loading control. 30nM of Py-S2 protein was sufficient to eliminate chromosomal SUMOylation, which is the equivalent concentration of endogenous PIASy protein in XEE, suggesting that the Py-S2 effectively deSUMOylates SUMOylated chromosomal proteins at a physiologically relevant concentration. Note that the concentration of dnUbc9 required for complete inhibition of chromosomal SUMOylation is 5μM in XEE, which is not within the physiological range and is difficult to induce a high expression level of dnUbc9 in cells. Addition of the Py-S2 C548A mutant (Py-S2 Mut) increased SUMO2/3 modification in chromosomal samples, including both TopoIIα SUMOylation and PARP1 SUMOylation. This suggests that the Py-S2 Mut acts as a dominant mutant for stabilizing SUMOylation.

**Supplemental Figure S2. Construction of Py-S2 and Py-S2 Mut DLD-1 cell lines.**

**(A)** Experimental scheme to introduce inducible Py-S2 and Py-S2 Mut into hH11 locus of DLD-1 cells. Cells were transfected with a donor plasmid with homology arms directed to the CCR5 locus (CCR5-TetON3G-mNeonPyS2-PuroR) and two gRNAs to target CCR5 locus. For the screening of the transgene integrated clones, primers were designed to amplify the 5’ region (∼3kb) and 3’ region (∼3.26kb) of the integration site.

**(B)** After the selection using 1ug/mL Puromycin, 1 clone each per construct were further subjected to genomic PCR to confirm the integration of the transgene.

**(C)** The whole cell lysates obtained from the candidate clones were subjected to Western Blotting to confirm the inducible expression of Py-S2 and Py-S2 Mut proteins. Anti-PIASy antibodies were used to detect expression of fusion proteins (+Dox) or not (-Dox), anti-H2A antibodies were used as a loading control.

**Supplemental Figure S3. Construction of OsTIR1 expressing DLD-1 cell lines.**

**(A)** Experimental schematic for the establishment of OsTIR1 gene expressing DLD1 cell. RCC1-OsTIR1-Myc-P2A-Blasticidin donor plasmid, and two guide RNAs targeting the 3’ end of RCC1 were used to integrate the OsTIR1 gene into the RCC1 locus.

**(B)** After the selection with 2ug/mL Blasticidin, fourteen clones were isolated and subjected to genomic PCR utilizing primers that targeted the 5’ end of the construct (upper panel). Non-transfected DLD-1 cells were used as a negative control (DLD-1 NC). Clones #48, 50, 52 and 56 were further verified by genomic PCR using primers for 3’ ends of the construct.

**(C)** Among the positive clones identified in **B**, two clones were chosen to verify the protein expression by Western blotting. Whole cell lysates obtained from asynchronous cell population were subjected to Western blotting. Non-transfected DLD-1 whole cell lysate was used as a negative control (DLD-1 NC). An anti-Myc antibody was used to detect OsTIR1 protein and anti-β-actin was used as a loading control. Clone #50 (marked in red) was chosen to utilize for subsequent AID tagging for TopoIIα and PICH.

**Supplemental Figure S4. Construction of TopoIIα-AID cell line.**

**(A)** Experimental schematic of donor plasmid tagging the 5’ end of endogenous TopoIIα with AID. Cells were transfected with the donor plasmid together with two different guide RNAs.

**(B)** After selection with 400ug/mL hygromycin, resistant clones were isolated. Whole cell lysate was obtained from cells and the expression of the transgene was screened by Western blotting analysis. Representative Western blotting of clones is shown. An anti-Flag antibody was used to detect AID-Flag tagged TopoIIα (∼190kDa) in the 700 channel (red) and anti-TopoIIα antibodies were used to detect both AID-Flag tagged TopoIIα and untagged TopoIIα (∼160kDa) in the 800 channel (green). Anti-β-tubulin was used as a loading control.

**(C)** Genomic DNA from hygromycin resistant clones was extracted for PCR analysis using indicated primers shown in A. Representative result of PCR amplification was shown. Clones showing only 3kbp DNA fragment are homozygous AID integrated clones (#72, #79 and #80).

**(D)** The clone #79 was treated with auxin for 2, 4, and 6-hours, and evaluated the TopoIIα depletion by Western blotting. As a control, DLD-1 OsTIR1#50 parental cells were treated with auxin for 6 hours (DLD1 TIR1). Whole cell lysates were subjected to Western blotting analysis using indicated antibodies. Clone #79 was chosen for further analysis in the subsequent experiments showed in Figure 2.

**(E)** DLD-1 cells with endogenous TopoIIα tagged with an auxin inducible degron (AID) were synchronized in mitosis and treated with auxin 6 hours after Thymidine release. Cells were plated onto fibronectin coated coverslips and subsequently stained with anti-TopoIIα, anti-CENP-C, and DNA was labeled with DAPI. TopoIIα foci on mitotic chromosomes are completely eliminated with auxin treatment.

**Supplemental Figure S5. Construction of PICH-AID cell line.**

**(A)** Experimental schematic of donor plasmid used to tag the 5’ end of endogenous PICH locus with AID tag. Cells were transfected with PICH-mAID-3xFlag-P2A-Hygromycin donor and two different guide RNAs. After selection with 400ug/mL hygromycin clones were isolated, whole cell lysates were collected from asynchronous populations, and Western blotting was performed.

**(B)** Representative Western blot for hygromycin-resistant clone screening is shown. An anti-Flag antibody was used to detect AID-Flag tagged PICH (∼180kDa) in the 700 channel (colored red) and anti-PICH antibodies were used to detect both AID-Flag tagged PICH (∼180kDa) and untagged PICH (∼150kDa) in the 800 channel (colored green). Non-transfected DLD-1 TIR1#50 parental cell line (labeled DLD-1) was used as a negative control. Anti-β-tubulin was used as a loading control. Among thirteen samples analyzed, the clones which showed a single yellow PICH band were chosen for genomic PCR analysis (clones #1, 6 and 11).

**(C)** Genomic DNA was isolated and subjected to PCR using an F1 primer located upstream of the left homology arm and Hygro Rev PCR primer located within the insert. Non-transfected DLD-1 TIR#50 parental cell DNA was used as a control (DLD-1 NC).

**(D)** The clones 1 and 6 were tested for further depletion of PICH protein by auxin addition at 4, 6, and 20-hour time points. The non-transfected DLD-1 TIR1#50 parental cells were used as a control with either non-treated (TIR#50) or treated with auxin for 20 hours (TIR#50 +Aux 20 hours). The whole cell lysates were subjected to Western blotting analysis. Anti-PICH antibodies were used to detect PICH (∼150kDa) or PICH-AID (∼180kDa), anti-β-tubulin antibodies were used as a loading control. Clone #1 (marked in red) was chosen to utilize for subsequent experiments showed in Figure 5 and Figure S6.

**(E)** DLD-1 cells with endogenous PICH tagged with an auxin inducible degron (AID) were synchronized in mitosis and treated with DMSO or ICRF-193. Auxin was added 6 hours after Thymidine release. Mitotic cells obtained by shake-off were plated onto fibronectin coated coverslips and subsequently stained with indicated antibodies. DNA was labeled with DAPI. PICH foci on mitotic chromosomes were completely eliminated with auxin in both DMSO and ICRF-193 treated cells.

**Supplemental Figure S6. Construction of Tet-inducible PICH mCherry mutants.**

**(A)** Experimental schematic of donor plasmid used to introduce PICH mCherry mutants into the CCR5 safe harbor locus. Cells were transfected with PICH-mCherry-P2A-Zeocin donor and two different guide RNAs.

**(B)** After selection with 400ug/mL Zeocin clones were isolated, cells were treated with doxycycline and auxin for 14 hours, whole cell lysates were collected from asynchronous populations, and Western blotting was performed.

**(C)** Genomic DNA was isolated and subjected to PCR using a Sv40 F primer located within the insert and CCR5 Rev located outside of the right homology arm. Non-transfected DLD-1 TIR#50 parental cell DNA was used as a control (DLD-1 NC).

**(D)** DLD-1 cells with endogenous PICH tagged with an auxin inducible degron (AID) and PICH mCherry mutants introduced into the CCR5 locus were synchronized in mitosis and treated with auxin and doxycycline for 22hours. Mitotic cells obtained by shake-off were plated onto fibronectin coated coverslips and subsequently stained with DAPI to label DNA and mCherry to label PICH expressing cells.

Primers used for amplification of homology arms

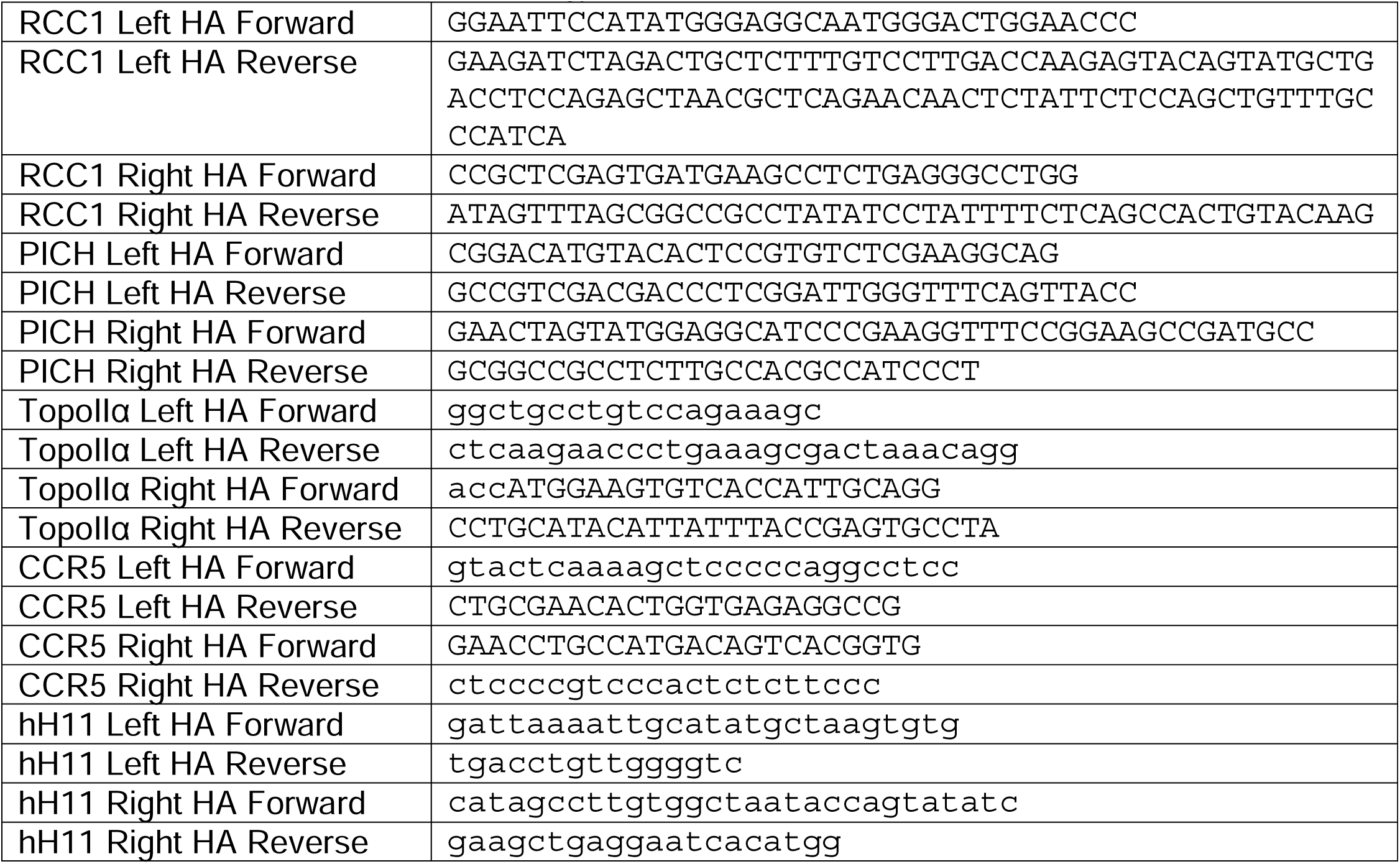

gRNA sequences used for Cas9 targeting of RCC1 locus or PICH locus

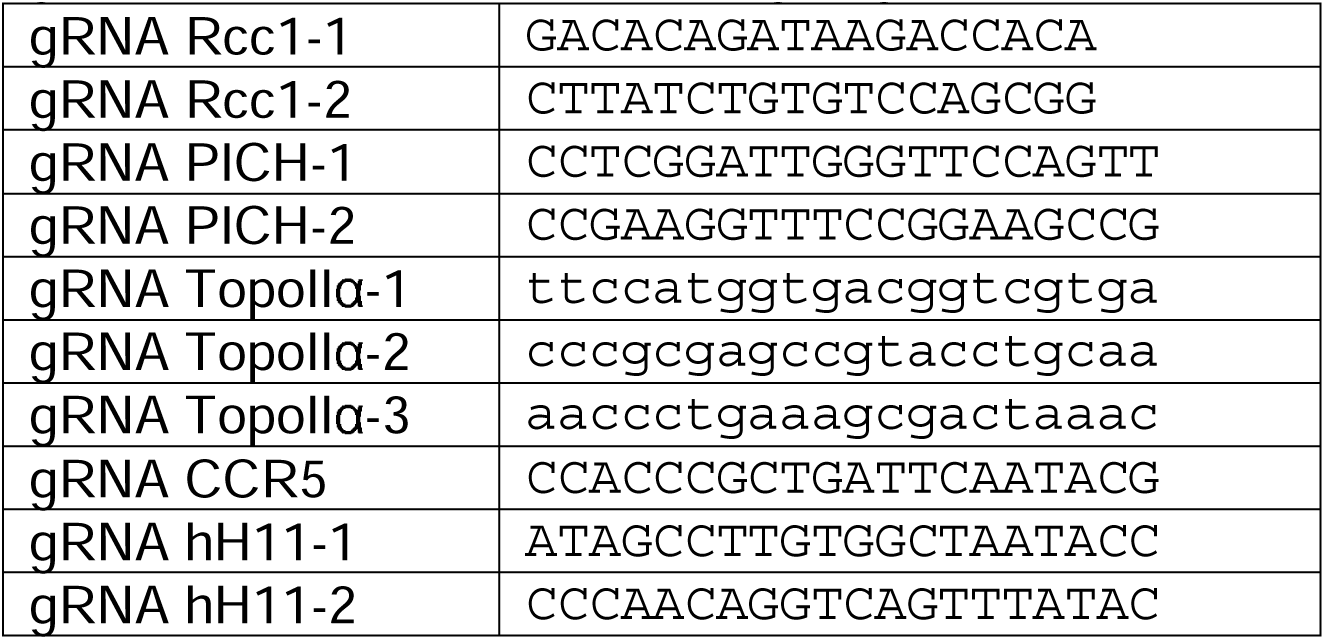

Primers used for genomic PCR

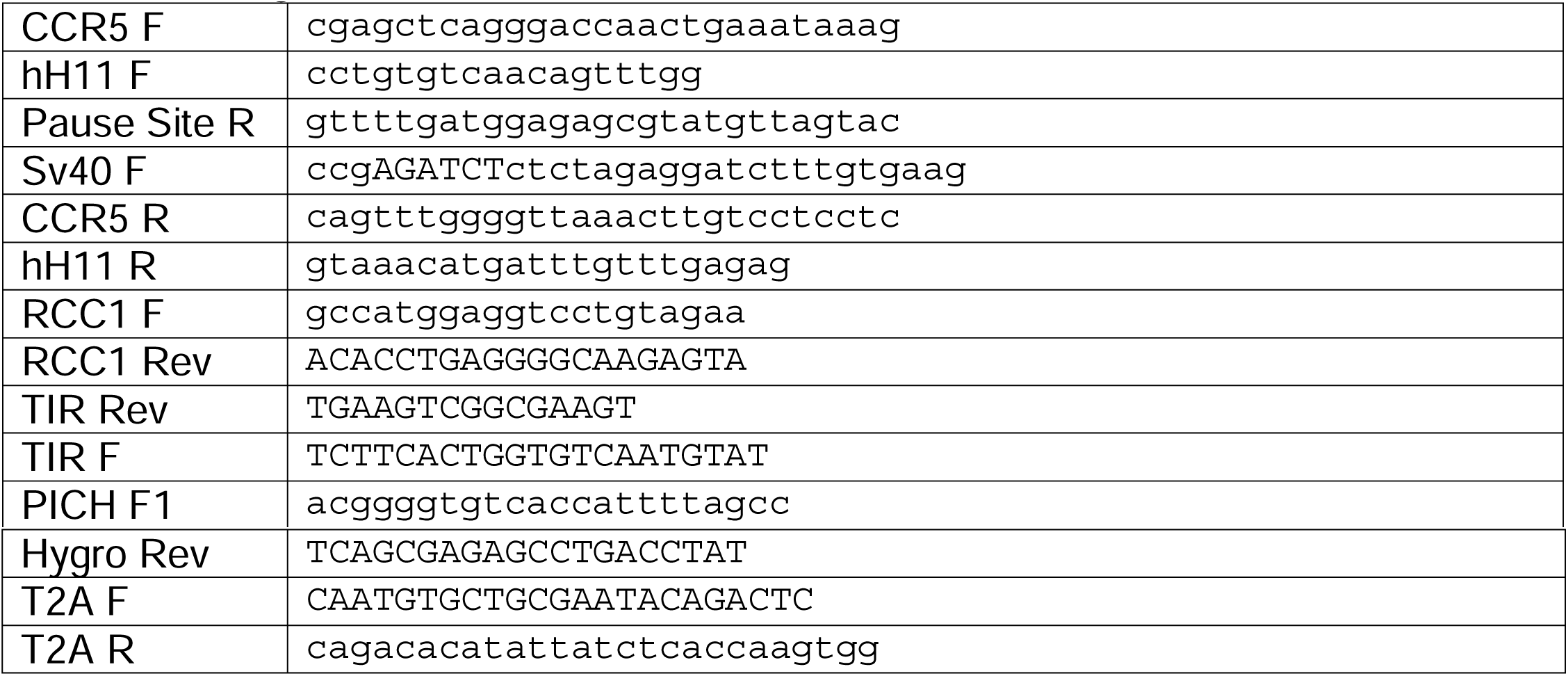

