## Supplemental figures for "PICH function is required for organization of SUMOylated proteins on mitotic chromosomes"

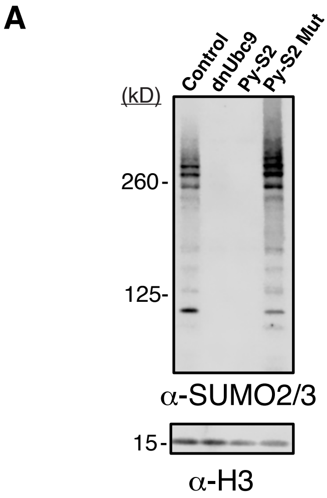


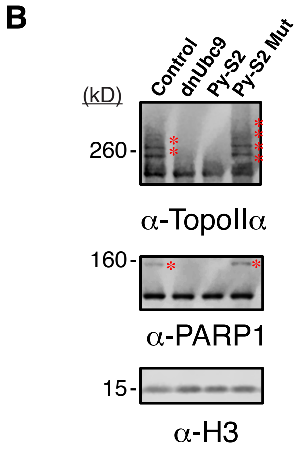


Supplemental Figure S1

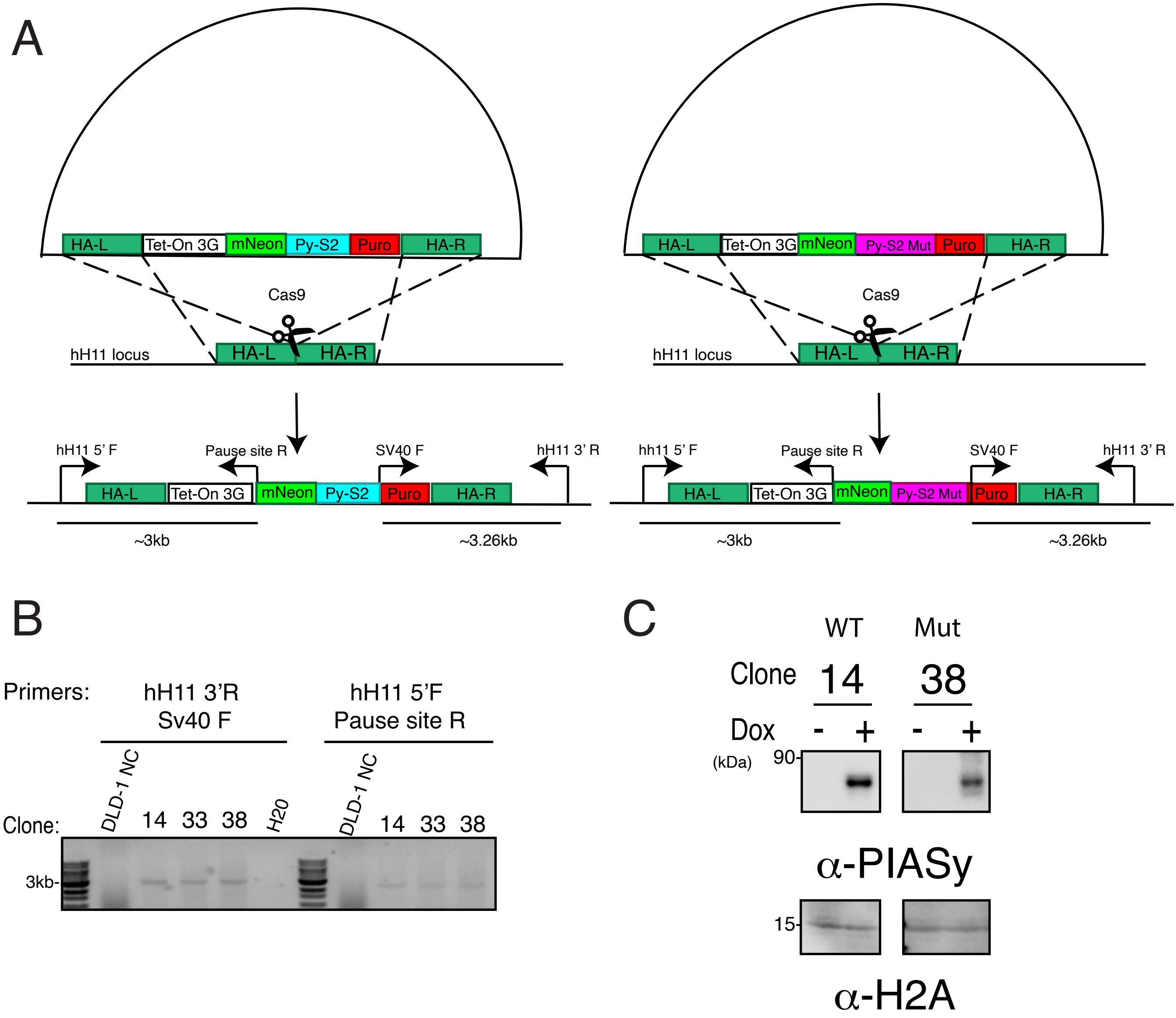


Supplemental Figure S2

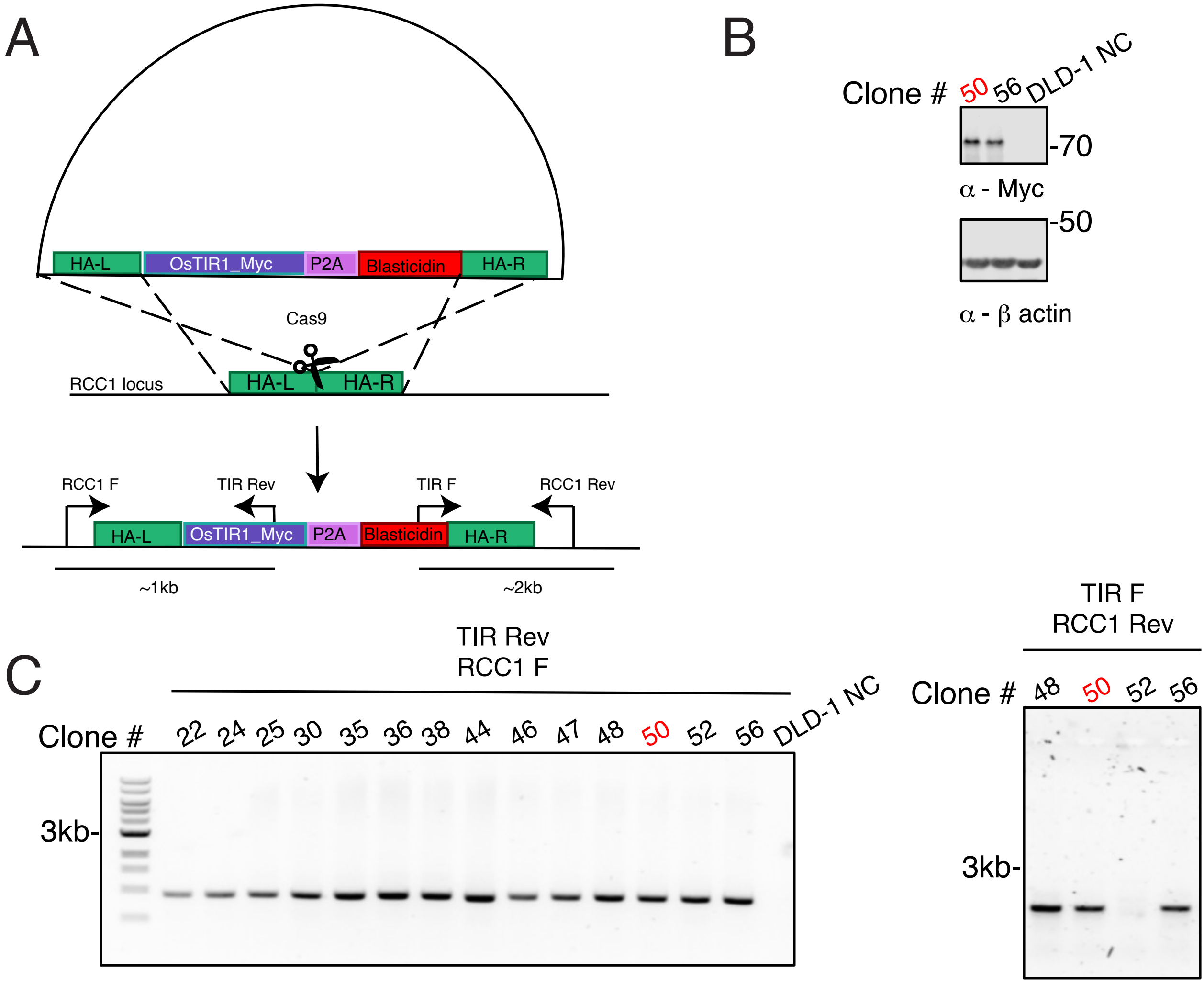


Supplemental Figure S3

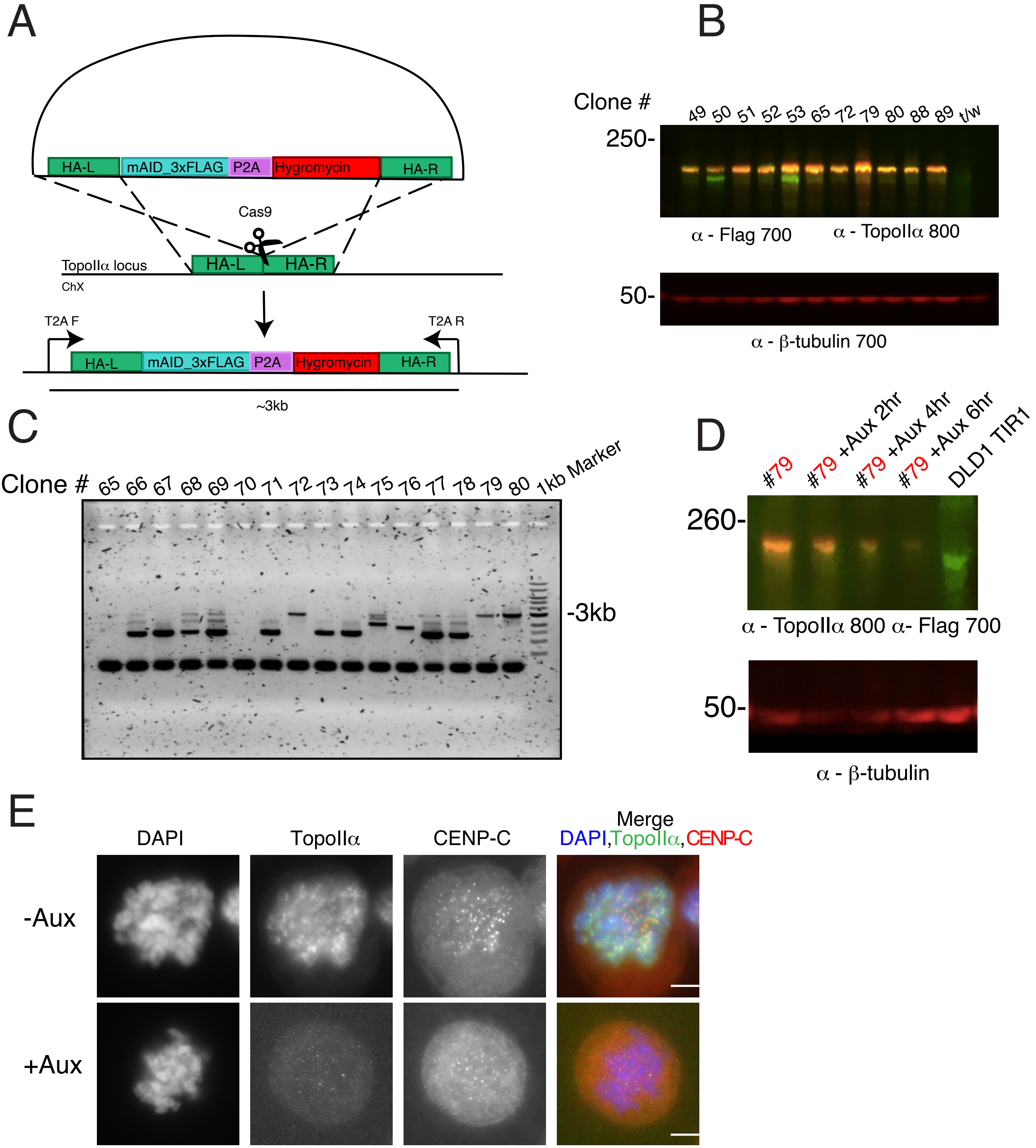


Supplemental Figure S4

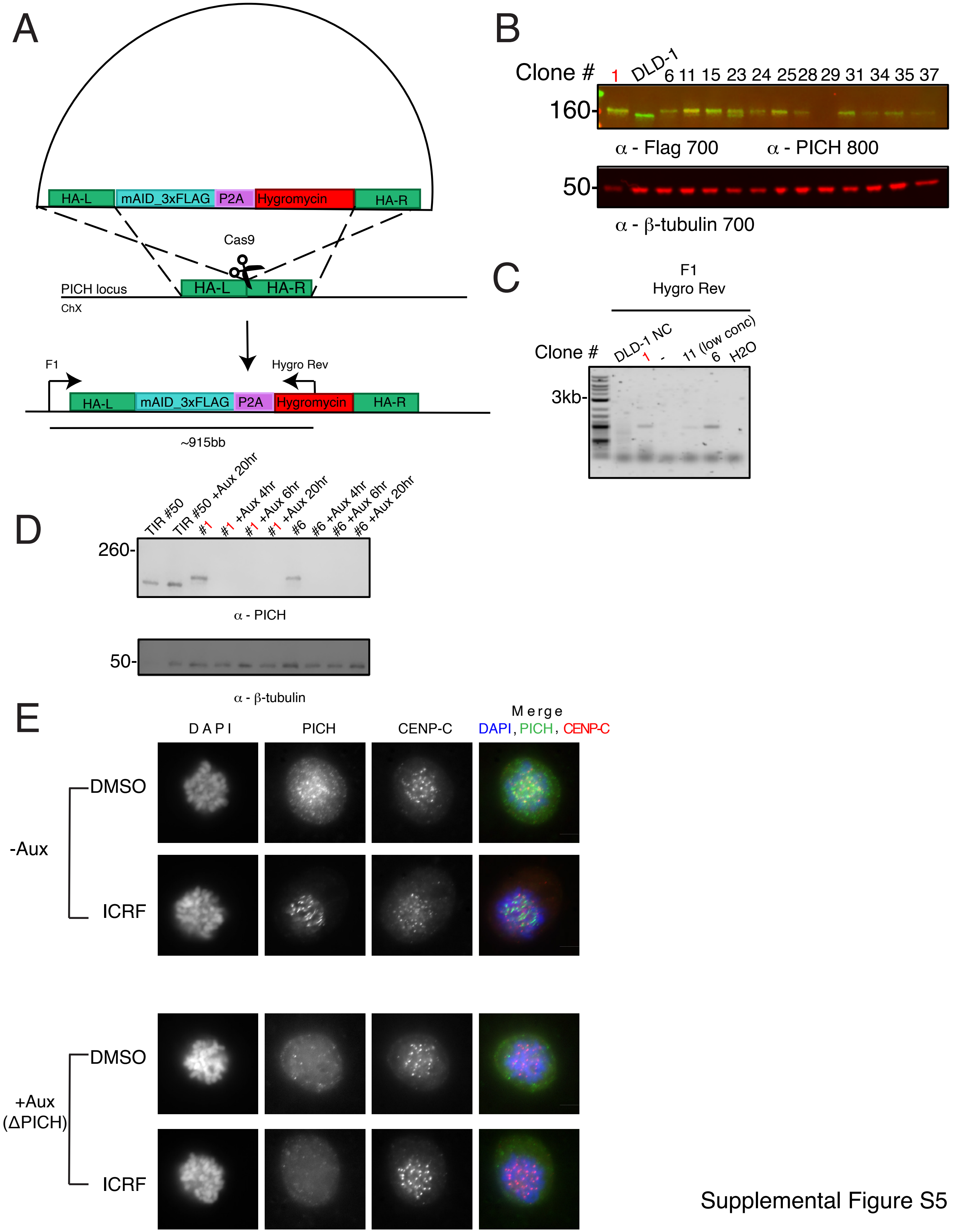


Supplemental Figure S5

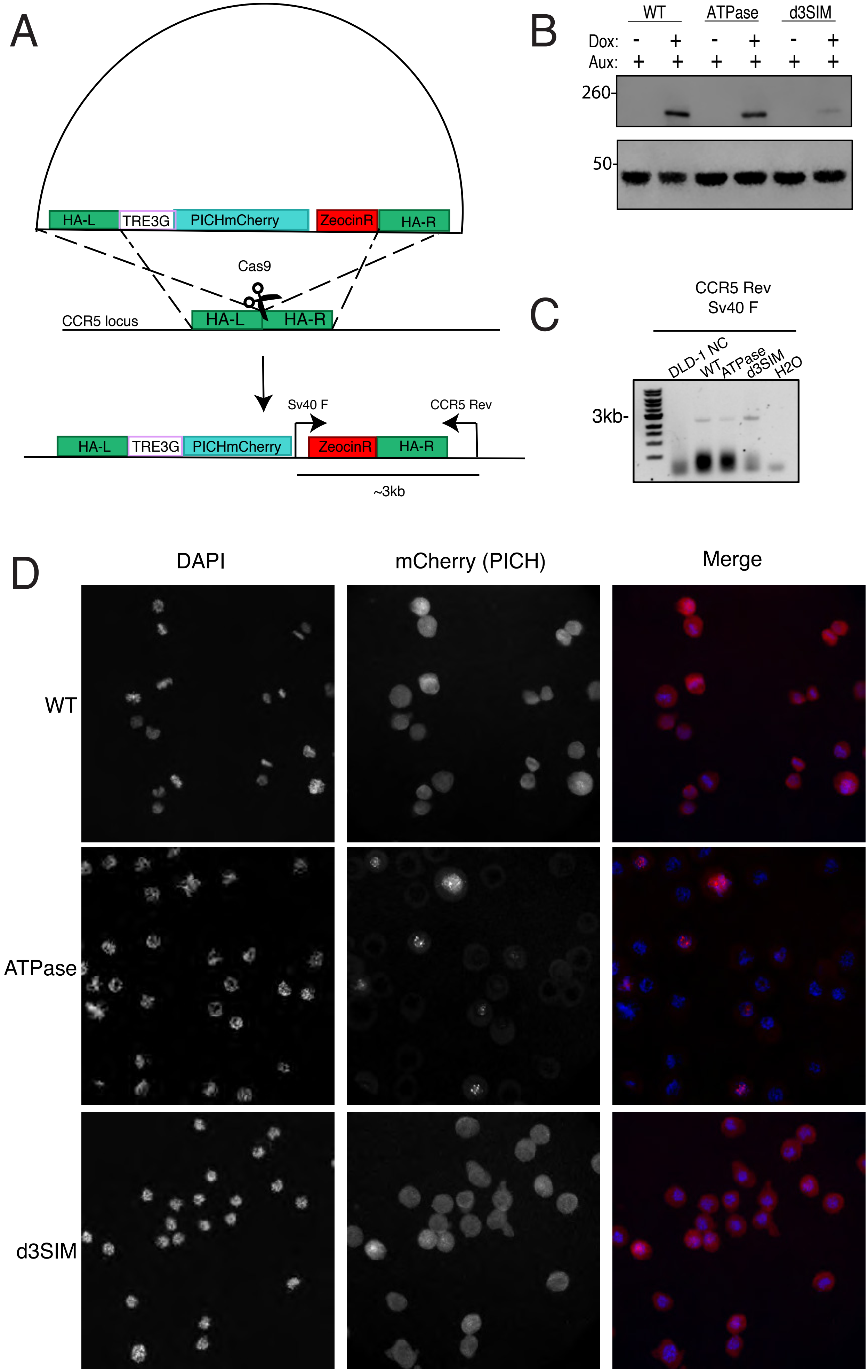


Supplemental Figure S6
